# An E-cadherin-actin clutch translates the mechanical force of cortical flow for cell-cell contact to inhibit epithelial cell locomotion

**DOI:** 10.1101/2021.07.28.454239

**Authors:** Ivar Noordstra, Mario Díez Hermoso, Lilian Schimmel, Alexis Bonfim-Melo, Denni Currin-Ross, Cao Nguyen Duong, Joseph Mathew Kalappurakkal, Richard G. Morris, Dietmar Vestweber, Satyajit Mayor, Emma Gordon, Pere Roca Cusachs, Alpha S. Yap

## Abstract

Adherens junctions allow cell contact to inhibit epithelial migration. But a long-standing puzzle is how locomotion is downregulated when E-cadherin adhesions form at surfaces perpendicular, but not those parallel, to the direction of migration. We now show that this arises from coupling between E-cadherin adhesions and the retrograde cortical flows of leader cells in migrating epithelia. At interfaces perpendicular to the direction of motion, such flows are antiparallel, which generates a tensile signal that induces the actin-binding domain of α-catenin to promote lateral growth of nascent adhesions and inhibit the lamellipodial activity necessary for migration. At interfaces parallel to the direction of motion, by contrast, cortical flows are aligned and no such mechanical inhibition takes place. Therefore, α-catenin mechanosensitivity in the clutch between E-cadherin and cortical F-actin allows cells to interpret the direction of motion via cortical flows and trigger the first signal for contact to inhibit locomotion.

## INTRODUCTION

Epithelial cells possess an intrinsic capacity to move that critically influences their tissue dynamics ^1^. Locomoting populations of cells are the foundation for the morphogenetically dynamic tissues seen in the early embryo ^2^ and in organs with high cellular turnover, like the gastrointestinal epithelium ^3^. But even apparently quiescent tissues retain a propensity to migrate that is rapidly activated during wound healing and tissue repair ^4^. Clearly, then, regulation of locomotility is a central factor in epithelial biology.

Here, a long-standing question is how contact between migrating cells regulates their movement ^5^. Many studies since the pioneering work of Abercrombie and Heaysman ^6, 7^ have documented the phenomenon of contact inhibition of locomotion (CIL) ^2, 8^. This has been implicated in tissue homeostasis and its loss proposed to contribute to disease, such as tumor invasiveness ^9^. In the classical description of CIL, migrating cells collapse their protrusions when they make contact, cease locomotion, then initiate new protrusions at free areas of their margins. This phenotype was first described in mesenchymal cells, such as cultured fibroblasts ^10^ and neural crest cells *in vivo* ^11^, but is also seen in epithelia ^12^. CIL has been identified in many different tissues, and a variety of intracellular signaling pathways have been implicated in its mechanism ^2^. However, less is known about the proximal steps that allow physical contact between the surfaces of cells to inhibit their locomotility.

In many cases, cell-cell contact is mediated by classical cadherin adhesion receptors. Cadherin adhesion begins with the homophilic *trans*-ligation of cadherin ectodomains presented on the surfaces of cells. This can then engage a variety of signaling pathways and regulators of the actin cytoskeleton, cellular processes with the potential to influence locomotility. So, it has been tantalizing to consider whether cadherins might also mediate CIL ^12^. Indeed, expression of classical cadherins conferred CIL behaviour on cadherin-null fibroblasts ^13^, while depletion of cadherins disrupted CIL in epithelia ^14^ and in the neural crest ^15^. These observations establish an important role for cadherins in CIL, but cadherin ligation by itself is not sufficient to explain how contact inhibits epithelial migration.

Of note, migrating epithelia are linked together by E-cadherin-based adherens junctions (AJ). Epithelial migration is inhibited when the leader cells at the front of migrating sheets encounter each other in a “head-on” fashion (Fig 1A). But elsewhere in the monolayer cells move despite forming cadherin adhesions with each other. This is clearly evident at the migrating front, where leader cells are connected to their neighbours by AJ at their sides (Fig 1A, side-side contacts). This implies that epithelial cells have a mechanism to evaluate the orientation of the cells with which they come into contact, discriminating head-on contacts that are perpendicular to the direction of movement from side-side contacts which are parallel to migration. This could reflect additional surface signals or, more parsimoniously, that cells detect some other important feature of cadherin adhesions found at head-on, but not at side-side, contacts. But what this feature might be is unknown.

**Figure 1.**
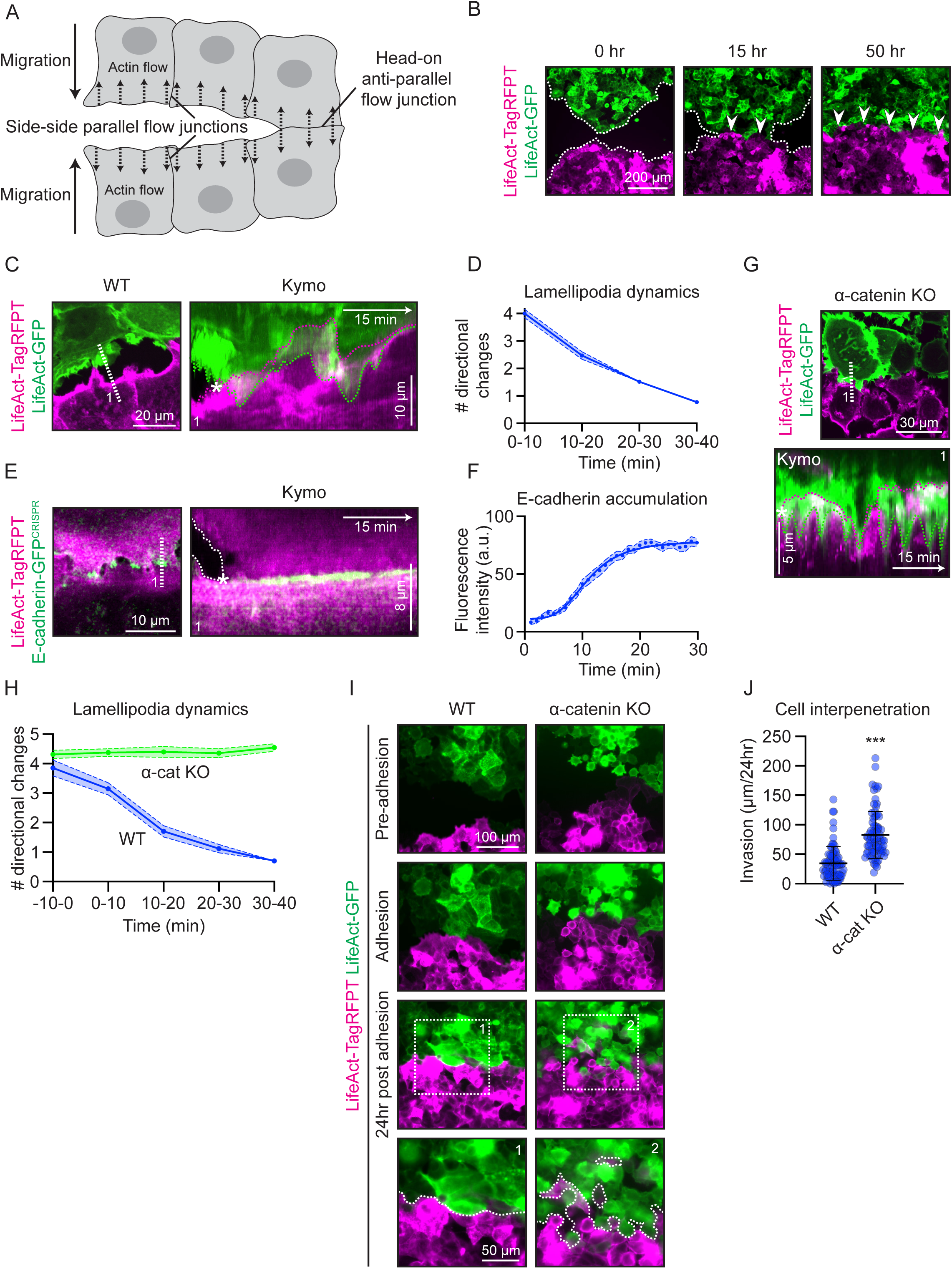
Nascent adherens junctions downregulate lamellipodia as cell-cell contact inhibits epithelial migration. **(A)** Schematic representation of contacts between epithelial cells in the migration assay. Left side: Migrating leader cells are connected to each other at side-side junctions where retrograde cortical flow is oriented in a parallel direction. Right side: migrating cells collide to make head-on contacts where retrograde cortical flows have an antiparallel orientation. Black solid arrows indicate direction of migration. Black dotted arrows indicate direction of actin flows. **(B)** Contact inhibition of migration results in a persistent, smooth boundary of contact when cell populations expressing either LifeAct-GFP (green) or LifeAct-TagRFPT (magenta) collide. Wavy dotted lines: free edges of the migrating populations; arrow heads: line of demarcation between former populations. Time points are measured after the first contact (0 hr). **(C, D)** Lamellipodial dynamics progressively decrease after contact between cells expressing LifeAct-GFP (green) or LifeAct-TagRFPT (magenta). (C) Representative image and kymograph from movie S1A. Wavy dotted lines: cell boundaries; white dotted line: site of kymograph. *First moment of contact. (D) Quantification of lamellipodia dynamics at overlapping contacts. Cell-cell contact events (n=71) from 3 independent experiments. Dots: means; light blue area: SEM. **(E, F)** Nascent adherens junction assembly between cells expressing E-cadherin-GFP^CRISPR^ (green) and LifeAct-TagRFPT (magenta). (E) Representative image and kymograph. Wavy dotted line: cell boundary; white dotted line: site of kymograph; *First moment of contact. (F) Quantification of E-cadherin intensity. Cell-cell contact events (n=34) from 4 independent experiments. Dots: means; light blue area: SEM; blue line: sigmoidal fit (R^2^=0.61). **(G, H)** Lamellipodial dynamics do not decrease after contact between α-catenin Knockout (KO) cells expressing LifeAct-GFP (green) or LifeAct-TagRFPT (magenta). (G) Representative image and kymograph. Wavy dotted lines: cell boundaries; white dotted line: site of kymograph. *First moment of contact. (H) Quantification of lamellipodia dynamics. Cell-cell contact events WT (n=34), α-cat KO (n=48) from 3 independent experiments. Dots: means; light colour areas: SEM. **(I, J)** Contact inhibition of migration is compromised by α-catenin Knockout as measured by interpenetration of cells beyond the boundary of contact between populations expressing either LifeAct-GFP (green) or LifeAct-TagRFPT (magenta). (I) Frames from representative movie S2A, B. White dotted lines: cell boundaries. (J) Quantification of cell interpenetration between WT or α-cat KO cell populations. Cell-cell contact events WT (n=80), α-cat KO (n=80) from 3 independent experiments. ***P<0,001; Mann-Whitney U test. Data are means ±SD with individual datapoints indicated.

From first principles, we might predict that any mechanism for contact to inhibit migration must act upon the locomotor apparatus of the moving cells. Lamellipodia are major organs of locomotility that are often apparent in migrating epithelial cells, particularly leader cells at the front of migrating sheets ^16^. Lamellipodia extend the cell margin forwards through branched actin assembly at their leading edges and retrograde flow of cortical actin, which generates force for protrusion when engaged with adhesions at the cell substrate ^17, 18^. Inhibition of actin assembly or retrograde flow compromises cell locomotion, supporting the notion that lamellipodia are important for cell migration. Now we report that cell-cell contact inhibits epithelial migration when newly-forming cadherin adhesions downregulate the lamellipodia that are necessary for cell locomotion. This involves a tension-activated clutch in cadherin-bound α-catenin that is activated when the antiparallel cortical flows found at head-on contacts become mechanically linked by cadherin adhesion.

## RESULTS

We used MCF7 mammary epithelial cells to study how contact regulates epithelial migration. Cells were grown to confluence in silicon moulds; migrated towards one another as coherent sheets when the moulds were removed; and formed a single monolayer connected by AJ upon contact with one another (Fig S1A). However, the line of demarcation between the former populations remained evident for many hours after they had established contact, as demonstrated by expressing differently-tagged LifeAct transgenes in the two populations. These revealed a smooth boundary between the two populations, with minimal interpenetration of cells (Fig 1B), suggesting that epithelial locomotion was inhibited when the migrating populations collided with one another.

### Nascent adherens junctions inhibit leader cell lamellipodia

We then focused on the leader cells to characterise how contact inhibited migration. Leader cells moved by extending large protrusive lamellipodia that were the first points of contact between the migrating populations (Fig 1C; movie S1A). Some cells separated almost immediately after this initial contact, whereas in other cells lamellipodia preserved overlapping contacts and became progressively less dynamic until they stopped after 30-40 min (Fig 1C, D, S1B; movie S1A). Using MCF7 cells whose endogenous E-cadherin had been tagged with GFP by CRISPR/Cas9 gene-editing (E-cadherin-GFP^CRISPR^), we found that the down-regulation of lamellipodial activity coincided with the appearance of immobile E-cadherin-GFP^CRISPR^ deposits (Fig 1E, F), ceasing shortly after E-cadherin levels had plateaued (Fig 1D, F). We refer to these E-cadherin deposits as nascent AJ, as they grew laterally to eventually formed the junctional border between the colliding cell populations (Fig S1A). This suggested that the assembly of nascent AJ might contribute to inhibiting lamellipodia.

To test this, we deleted α-catenin, a critical component of the cadherin adhesion complex, by CRISPR/Cas9 gene-editing (α-catenin KO) (Fig S1C-E). Confluent α-catenin KO cells expressed E-cadherin at their cell surface but did not assemble AJ (Fig S1E-F). Moreover, contact between α-catenin KO cells failed to inhibit lamellipodia in migration assays (Fig 1G,H; movie S1B). This was accompanied by a failure of contact to inhibit locomotion. α-catenin KO populations marked with differently-tagged LifeActs interpenetrated after contact, forming finger-like projections rather than the smooth borders seen with WT cells (Fig 1I, J; movie S2). These findings reinforce the notion that nascent AJ allow contact to inhibit epithelial locomotion. They further suggest that a key might lie in downregulating the lamellipodial activity that is necessary for leader cells to migrate.

### Adherent and free surface cadherins couple to cortical flow in migrating leader cells

Lamellipodia are distinguished by retrograde flows of cortical actin that generate shear forces for cell protrusion ^18^. Therefore, to understand how assembly of AJ might inhibit lamellipodia, we analysed the relationship between E-cadherin and cortical actin flow in the leader cells. Leader cells were distinguished by prominent retrograde cortical flows that were evident both in the free lamellipodia and in the side-to-side contacts that linked leader cells together (Fig 2A; movie S3A). As seen elsewhere ^19^, E-cadherin adhesions at the side-to-side contacts displayed retrograde movement, consistent with their being coupled to the flowing cortex (Fig 2B; movie S3B).

**Figure 2.**
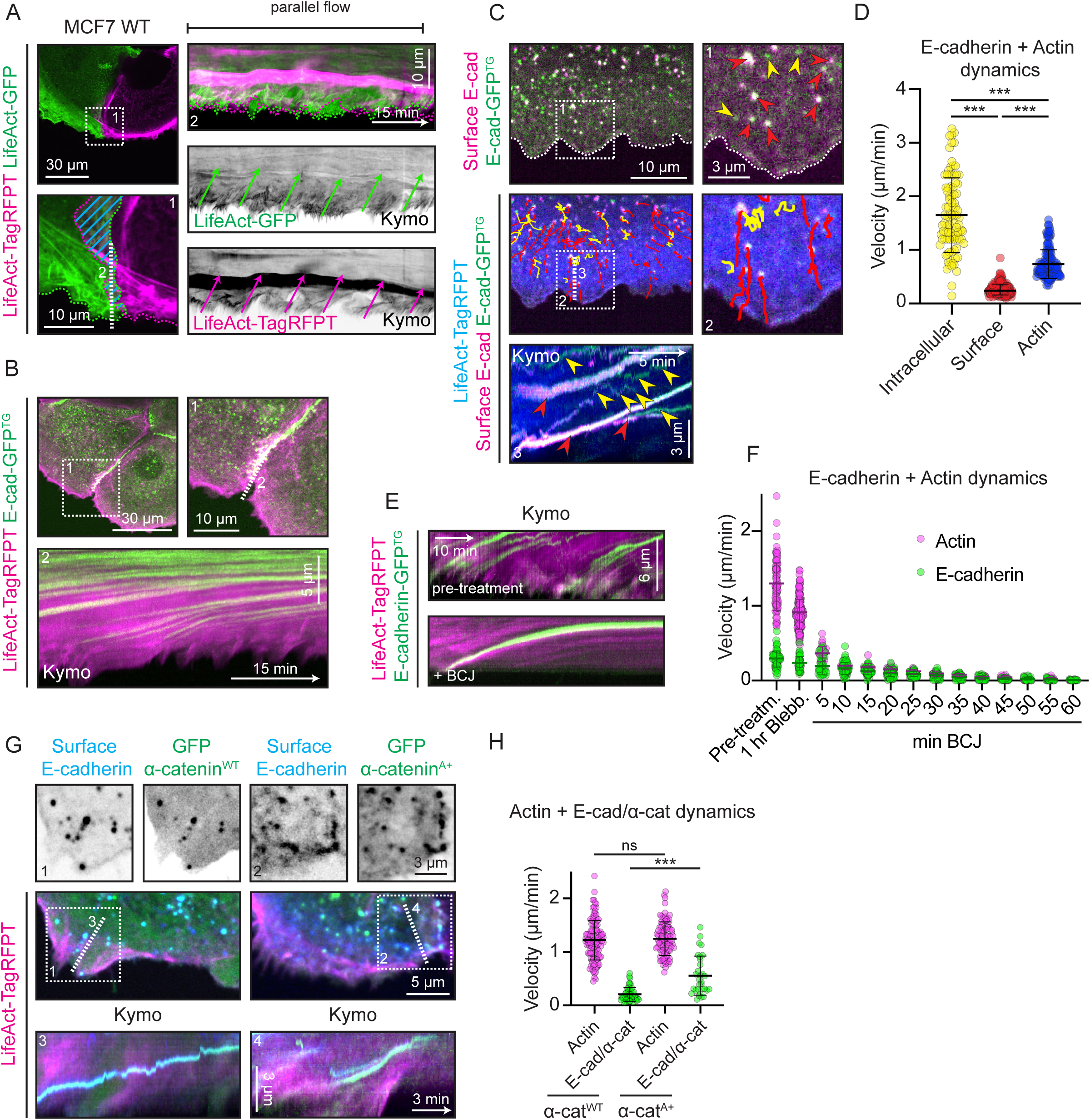
Free and adherent E-cadherin is coupled to retrograde flow in migrating leader cells. **(A)** Parallel actin flow at side-side junctions between leader cells expressing LifeAct-GFP (green) or LifeAct-TagRFPT (magenta). Representative image and kymograph from movie S3A. Wavy dotted lines: cell boundaries; blue lines: overlapping area; green and magenta arrows: direction of actin flows; white dotted line: site of kymograph. **(B)** E-cadherin-GFP^TG^ (green) at side-side contacts displays retrograde flow with the parallel actin flows marked by LifeAct-TagRFPT (magenta). Representative image and kymograph from movie S3B. White dotted line: site of kymograph. **(C, D)** E-cadherin is found as surface clusters and intracellular vesicles in the lamellipodia of migrating leader cells expressing LifeAct-TagRFPT (blue) and E-cadherin-GFP^TG^ (green). Surface E-cadherin is labelled in live cells with extracellular E-cadherin mAbs (magenta). E-cadherin-GFP^TG^ puncta that are not surface-labelled appear green while puncta that are surface-labelled appear white. (C) Representative image and kymograph from movie S4. Wavy white dotted line: free cell edge; straight white dotted line: site of kymograph; yellow lines and arrow heads: E-cadherin-containing intracellular vesicles; red lines and arrow heads: E-cadherin surface clusters. (D) Quantification of E-cadherin motility and actin flow rates. Intracellular vesicles (n=94), Surface clusters (n=167), Actin flow waves (n=136) from 3 independent experiments. **(E, F)** Effect of Blebbistatin, CK666, Jasplakinolide (BCJ) treatment on retrograde flow of surface E-cadherin clusters (green) and F-actin (magenta). (E) Representative kymographs. Site of kymograph is marked as a white dotted line in Fig S2E. (F) Quantification of surface E-cadherin clusters and F-actin flow rates. Actin flow waves (n=410), surface E-cadherin clusters (n=554) from 3 independent experiments. **(G, H)** Increasing the affinity of α-catenin for F-actin increasing the coupling of surface E-cadherin clusters to retrograde cortical flow. Comparison of actin flow and surface E-cadherin cluster movement in leader cells expressing LifeAct-TagRFPT (magenta) and GFP-α-catenin^WT^ or GFP-α-catenin^A+^ (green). Surface E-cadherin is labelled with surface-delivered E-cadherin mAbs (blue). (G) Representative images and kymographs. White dotted lines: site of kymographs. (H) Quantification of actin flow and flow of surface-labelled E-cadherin clusters that contained GFP-α-catenin transgenes (E-cad/α-cat). Actin (α-cat^WT^) flow waves (n=117), E-cad/α-cat (α-cat^WT^) clusters (n=49), Actin (α-cat^A+^) flow waves (n=95), E-cad/α-cat (α-cat^A+^) clusters (n=31) from 2 independent experiments. ns: not significant, ***P<0,001; Kruskal-Wallis test. Data are means ±SD with individual datapoints indicated.

Puncta of E-cadherin were also evident on closer inspection of the free lamellipodia, as visualized with either E-cadherin-GFP^CRISPR^ (Fig S2A) or with an exogenously-expressed transgene (E-cadherin-GFP^TG^) (Fig 2C). The greater signal strength of E-cadherin-GFP^TG^ revealed that the puncta comprised two subpopulations with distinct patterns of movement. One population showed rapid (~1.5 µm/min) movements in both anterograde and retrograde directions (Fig 2C, D, yellow; movie S4); often appeared to move along microtubules (Fig S2B); and were not accessible to extracellular E-cadherin mAbs in live cells (Fig 2C, yellow). We consider these to be E-cadherin-containing intracellular vesicles.

In contrast, ~50% of E-cadherin-GFP^TG^ puncta formed when diffuse cadherin coalesced at the leading edges of the lamellipodia (Fig S2C). These showed a distinctive, slow processive retrograde movement (~0.2 µm/min) (Fig 2C, D, red; movie S4) that made them readily distinguishable from the intracellular vesicles. They co-labelled in live cells with extracellular E-cadherin mAbs (Fig 2C, red), but were not due to mAb-induced cross-linking, because their size was unaffected by the mAbs. Since they immunostained for β- and α-catenin (Fig S2D), we conclude that they represent *cis*-clusters of E-cadherin-catenin complexes that form independently of adhesive *trans-*ligation on the free cell surface, as previously reported ^20, 21^. (We refer to these as “free surface” or “unligated” cadherin clusters to emphasize that they do not represent *trans*-ligated complexes.)

Two observations indicated that the retrograde flow of free surface cadherin clusters reflected their association with cortical flow. First, the retrograde flow of surface cadherin clusters was inhibited by a cocktail of blebbistatin, CK666, and jasplakinolide that effectively suppressed actin flow (Fig 2E, F, S2E). Second, unligated cadherin clusters moved more slowly than cortical actin flow, but this was accelerated if their affinity for F-actin was increased. This was evident when we expressed a transgene that is mutated to unfold the actin-binding domain (ABD) and increase its affinity for F-actin (α-catenin^A+^ (Fig S3A, B); ^22^). Surface E-cadherin moved faster in cells expressing α-catenin^A+^ than in controls, although the speed of cortical flow was not itself affected (Fig 2G, H). By implication, α-catenin coupled free surface cadherins to the flowing cortical cytoskeleton and, indeed, controlled the efficiency of coupling via its affinity with actin filaments. (We could not test the effect of α-catenin KO as this reduced free cadherin surface clustering (Fig S2F, G), making it difficult to measure cadherin movement.) Therefore, in leader cells both free cadherins on lamellipodia and cadherins at side-to-side contacts were coupled to retrograde cortical flow.

### Coupling antiparallel cortical flows allows E-cadherin adhesion to inhibit lamellipodia

The distinct movements of surface versus vesicular E-cadherin allowed us to define how nascent AJ assembled. Nascent AJ grew when slow moving free surface cadherin clusters incorporated into the immobile E-cadherin deposits that represent nascent AJ, often fusing with the ends, leading to lateral extension of the AJ (Fig 3A). But we did not detect incorporation of rapidly-moving puncta that would represent intracellular vesicles. This implied that nascent AJ assembled when free E-cadherin clusters that were coupled to cortical flow underwent *trans*-ligation as lamellipodia made contact with each other. Since E-cadherin in the nascent AJ was immobile, we then asked if contact affected retrograde flow in lamellipodia.

**Figure 3.**
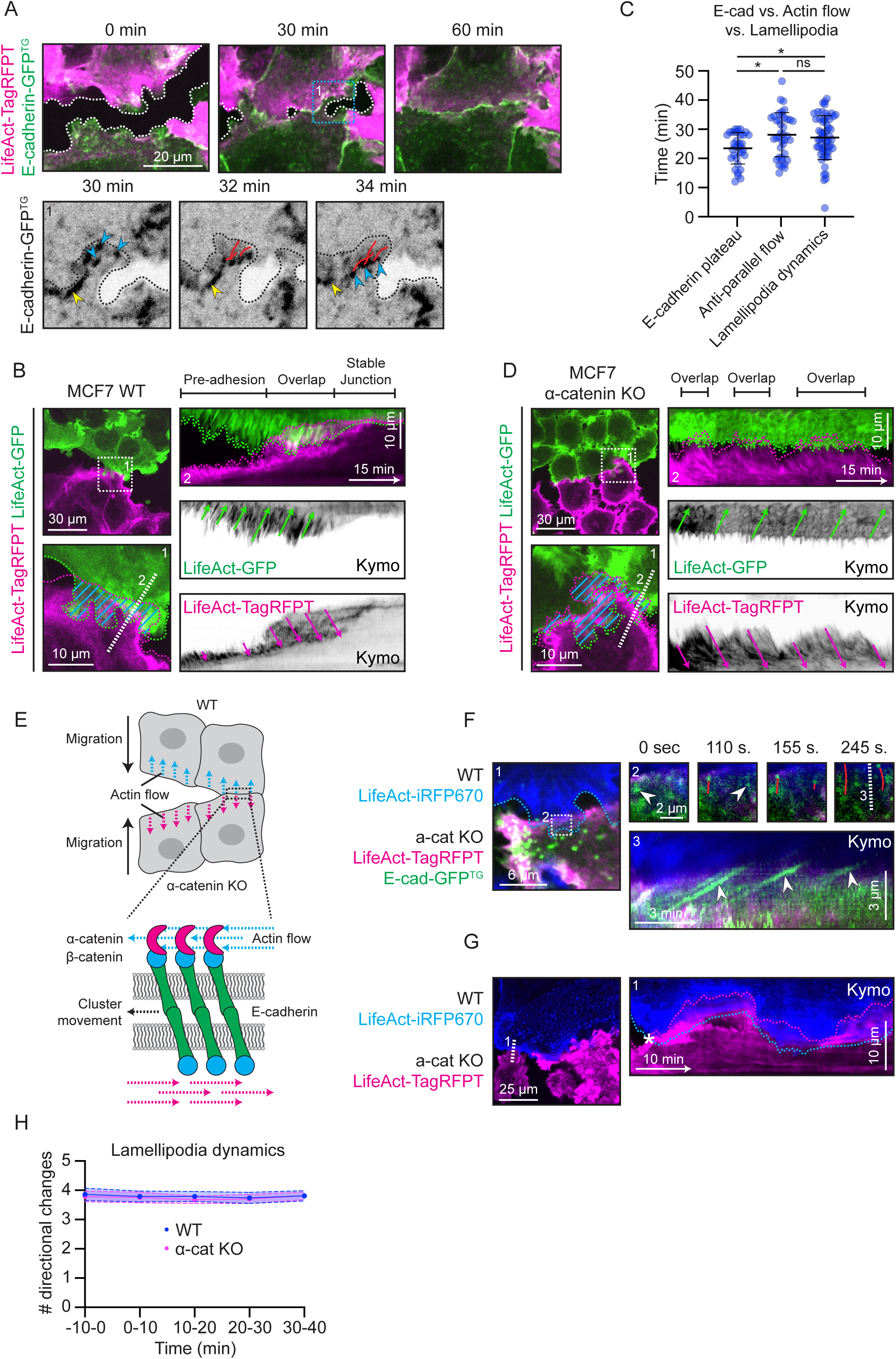
Nascent adherens junctions couple antiparallel cortical flows together when contact inhibits locomotion. **(A)** Nascent adherens junction assembly on contact between migrating populations expressing LifeAct-TagRFPT (magenta) and E-cadherin-GFP^TG^ (green): Timelapse overview (top) and magnified detail (from 1, below). Movement pattern of E-cadherin-GFP^TG^ clusters (blue arrowheads) that incorporated into already-static E-cadherin accumulations representing nascent AJ (yellow arrowheads). Tracks of cluster movement (red lines) are consistent with the slow processive movement of surface clusters. Black dotted lines: cell edge of bottom cell. **(B)** F-actin flows are anti-parallel at head-on junctions between WT cells expressing LifeAct-GFP (green) or LifeAct-TagRFPT (magenta). Wavy dotted lines: cell boundaries; blue lines: overlapping area; green and magenta arrows: direction of actin flows; white dotted line: site of kymograph. **(C)** E-cadherin accumulation in nascent AJ plateaus before antiparallel actin flows and lamellipodia cease. Cell-cell contact events for E-cadherin (n=34) from 4 independent experiments, actin flow (n=39) from 3 independent experiments, lamellipodia (n=58) from 3 independent experiments. **(D)** Anti-parallel actin flow at head-on contacts between migrating α-catenin KO cells expressing LifeAct-GFP (green) or LifeAct-TagRFPT (magenta). Representative image and kymograph from movie S1B. Wavy dotted lines: cell boundaries; blue lines: overlapping area; green and magenta arrows: direction of actin flows; white dotted line: site of kymograph. **(E)** Schematic representation of heterologous contacts between WT and α-catenin KO cells in migration assays. Note that the cadherin/catenin complexes are coupled to cortical actin flows in WT cells but not in α-catenin KO cells. Black arrows indicate direction of migration. Colour arrows indicate direction of actin flows. Black dotted arrow indicates direction of cadherin/catenin cluster movement. **(F)** Heterologous contacts between migrating WT cells (expressing only LifeAct-iRFP670, blue), and α-catenin KO cells (expressing both LifeAct-TagRFPT, magenta; and E-cadherin-GFP^TG^, green) as in (E). Representative image and kymograph from movie S5. Note that E-cadherin is only marked in the α-catenin KO cells. Blue dotted line: WT cell edge; white dotted line: site of kymograph; white arrow heads: anterograde movement (Kymo) of E-cadherin clusters in α-catenin KO cells; red lines: tracks of moving clusters. **(G, H)** Lamellipodial dynamics do not decrease after heterologous contact between WT cells expressing LifeAct-iRFP670 (blue), and α-catenin KO cells expressing LifeAct-TagRFPT (magenta) as in (E). (G) Representative image and kymograph of the zone of overlap at contact. Wavy dotted lines: cell boundaries; white dotted line: site of kymograph. *First moment of contact. (H) Quantification of lamellipodia dynamics in both the WT and α-catenin KO cells at overlapping heterologous contacts. Cell-cell contact events (n=28) from 2 independent experiments. Dots: means; light colour area: SEM. ns: not significant, *P<0,1; Kruskal-Wallis test. Data are means ±SD with individual datapoints indicated.

To do this, we examined kymographs of LifeAct tagged with different fluorophores in the colliding populations. As expected from the topology of the migrating cells (Fig 1A), cortical flows were oriented in opposite (antiparallel) directions when migrating lamellipodia made head-on contacts with one another (Fig 3B; movie S1A). This contrasted with the side-side junctions between migrating leader cells, where flows were oriented in the same (parallel) direction (Fig 1A, 2A; movie S3A). Interestingly, these oppositely-directly cortical flows persisted at the overlapping contacts until their lamellipodia became quiescent (Fig 3C; movie S1A). Both opposed retrograde flows and lamellipodial activity ceased ~ 5-10 min after the peak of nascent AJ assembly (Fig 3C). This suggested that AJ assembly might inhibit lamellipodia by downregulating their underlying retrograde flows. One potential mechanism could involve nascent AJs linking the oppositely-directly flows together. Because free cadherins already engaged the cortex, they would be predicted to physically couple flows together when they underwent adhesive *trans-*ligation. In support of this idea, we found that cell-cell contact failed to inhibit retrograde flow when E-cadherin was uncoupled from the cortex in α-catenin KO cells (Fig 3D; movie S1B). However, as α-catenin KO also compromises cadherin clustering (S2F, G), it remained possible that some other aspect of cadherin signaling inhibited cortical flow.

To pursue this hypothesis, we studied heterologous cell-cell contacts, where E-cadherin was uncoupled from the cortex in only one cell of a contact (Fig 3E). We reasoned that if coupling of antiparallel retrograde flows allows cadherin to inhibit lamellipodia, then it would be necessary for *trans-*ligated cadherin complexes to link to the cortex on *both* sides of a nascent adhesion. So, we analysed the head-on collisions that were made when separate populations of WT and α-catenin KO cells migrate towards one another. Of note, E-cadherin clustering was restored to the α-catenin KO cells at these heterologous contacts (Fig 3F; movie S5). We interpret this to reflect *trans-*ligation between cadherin in the KO cells with E-cadherin clusters formed in the WT cells, evidence that cadherins were functional in the α-catenin KO cells. Furthermore, cadherin clusters in the α-catenin KO cells also demonstrated slow processive movement; but this occurred in an anterograde direction, implying that they were being pulled by retrograde flow in the WT cells (Fig 3F; movie S5). This confirmed that cortical binding of cadherins was compromised in the α-catenin KO cells. Therefore, this set up preserved cadherin clustering and connection to flow, but disrupted the connection between flows across the contacting membranes that is expected when the cadherin adhesion system is intact.

Nonetheless, heterologous contacts failed to down-regulate either retrograde flows or the lamellipodial activity that they drive (Fig 3G, H). This reinforces the concept that mechanical coupling of anti-parallel cortical flows is a key that allows nascent AJ to inhibit lamellipodial activity at head-on contacts. Of note, retrograde flow and lamellipodial activity continued unabated by contact in the WT cells, as well as the α-catenin KO cells (Fig 3G, H). This cell-nonautonomous behaviour strengthens the notion that coupling of retrograde flows between both colliding cells was necessary for their down-regulation by nascent AJ.

### Mechanical tension across cadherin adhesions correlates with the topology of cortical flows

But how might mechanical coupling of cortices allow AJ to inhibit retrograde flows at head-on contacts, but not at the side-to-side AJ that connected migrating leader cells? An important consideration lay in how the different topologies of flow might affect mechanical tension applied to the cadherins. Cortical flows can exert mechanical force on membrane proteins ^23, 24^. We confirmed this for *trans*-ligated cadherins by plating MCF7 cells onto soft PDMS substrata coated with E-cad ectodomain ligands (Fig 4A, Fig S4A). Under these conditions, cellular E-cadherin was immobilized and retrograde cortical F-actin flow persisted at the cell-substrate interface in the cortical regions of the cells (Fig 4B). Microbeads embedded in the substrata underwent a periodic displacement that coincided exactly with waves of F-actin passing over the beads (Fig 4C, D), evidence that force applied from cortical flow was transmitted across *trans*-ligated cadherin complexes. Based on this test of principle, we then predicted that the tug-of-war between antiparallel flows at head-on contacts would generate greater tension than the parallel flows found at side-side contacts (Fig 1A).

**Figure 4.**
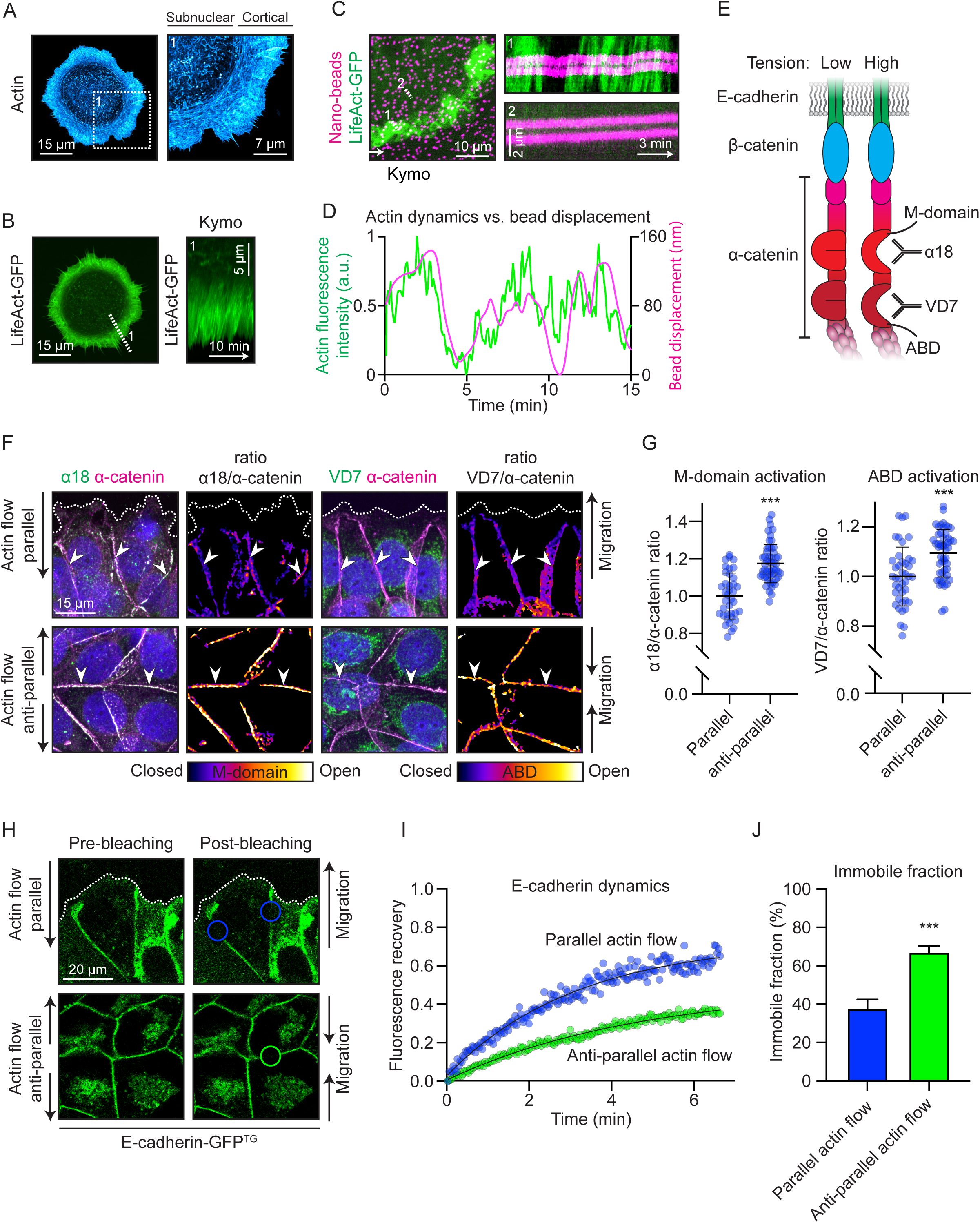
Cortical flow exerts mechanical tension on *trans*-ligated E-cadherin complexes. **(A)** F-actin organization in MCF7 cell spread on E-cadherin ectodomain-coated substrate and stained with phalloidin. **(B)** F-actin (LifeAct-GFP) dynamics in the cortical region of an MCF7 cell spread on E-cadherin ectodomain-coated substrate. White dotted line: site of kymograph. **(C, D)** Transmission of actin flow forces across *trans*-ligated cadherin complexes. (C) Cortical zone of a cell expressing LifeAct-GFP (green), spread on an E-cadherin-coated PDMS substrate (2-3 kPa) containing nanobeads (magenta). White dotted lines: site of kymographs. (D) Quantification of actin dynamics and bead displacement in kymograph from C1. **(E)** Schematic representation of the cadherin/catenin complex under low and high tension. Tension-sensitive α-catenin domains and antibodies (α18 and VD7) that report the conformational changes are indicated. **(F, G)** Tension-sensitive conformational changes in α-catenin at side-side vs head-on junctions between leader cells. (F) Representative images of side-side (top row) and head-on (bottom row) junctions stained for α18 (M-domain) or VD7 (ABD) (green) and total α-catenin (magenta). Ratiometric α18/α-catenin and VD7/α-catenin images indicate domain-specific conformational changes. White arrowheads indicate examples of analysed junctions. (G) Quantification of α-catenin M-domain (α18) and ABD (VD7) conformational changes at parallel (side-side) and anti-parallel (head-on) flow junctions. Parallel flow junctions: M-domain (n=38) or ABD (n=41); anti-parallel flow junctions M-domain (n=55) or ABD (n=52) from 3 independent experiments. **(H, I, J)** E-cadherin stability measured by fluorescence recovery after photobleaching in side-side vs head-on junctions. (H) Representative images of Pre- and Post-bleaching areas at parallel (side-side) and anti-parallel (head-on) flow junctions in cells expressing E-cadherin-GFP^TG^. (I) Normalized fluorescence intensity graphs after photobleaching. Parallel flow junctions (n=20), anti-parallel flow junctions (n=9) from 2 independent experiments. Dots: means; Black lines: logistic fits (Parallel actin flow, R^2^=0.97; Anti-parallel actin flow, R^2^=0.98). (J) E-cadherin-GFP^TG^ immobile fractions after photobleaching. ***P<0,00l; Mann-Whitney U test. Data are means ±SD with individual data points indicated (G) or means +SEM (J).

To test this, we probed for tension-sensitive changes in the conformation of α-catenin at head-on versus side-side contacts. Mechanical tension opens the central M-domain of α-catenin ^25^ and also unfolds the autoinhibited alpha1 helix in the F-actin-binding site (ABD) ^22^; these conformational changes are detectable with the α18 monoclonal ^25^ and VD7 polyclonal antibodies ^26^, respectively (Fig 4E). Consistently, we found that staining for both α18 and VD7, corrected for total α-catenin, was more intense in AJ at head-on contacts compared with side-to-side contacts (Fig 4F, G). Therefore, tension across the cadherin complex was higher at the head-on contacts where AJ inhibited lamellipodia, than at the side-side contacts. Of note, we did not detect cortical Myosin II at these early contacts (not shown), suggesting that actin flow might be the direct source of mechanical force. This was further supported by differences in E-cadherin stability at these two contacts. After photobleaching, E-cadherin-GFP fluorescence recovered more slowly at head-on contacts than at side-side contacts (Fig 4H-J). As tension stabilizes E-cadherin ^27, 28^ this difference in cadherin stability was consistent with greater tension at head-on contacts.

### Modelling cadherin-actin interactions as a tension-activated clutch

We then considered how greater tension across the cadherin complexes at head-on contacts could inhibit retrograde flows and lamellipodial activity. One possibility was if this tensile stimulus increased the mechanical coupling of anti-parallel flows by nascent AJ. The tension-sensitive conformational changes in α-catenin that we observed at head-on contacts increase its association with the actin cytoskeleton ^29^. Opening of the ABD increases its affinity for F-actin to mediate a catch-bond ^22, 30^; and opening of the M-domain recruits vinculin, an additional F-actin-binding protein ^25^. We focused on the effect of tension on the ABD, since a role for the M-domain was not evident in screening studies where an α-catenin mutant that cannot recruit vinculin (a-catenin^V-^ (S3A, B)) fully restored AJ to α-catenin KO cells (Fig S5A).

We then considered a simple model of mechanical antagonism, where *trans-*ligated cadherins at head-on contacts would allow antiparallel flows to pull against one another, causing their reciprocal inhibition (Fig 5A). Decreased cortical flow would downregulate lamellipodial activity to reduce locomotility, as well as decrease the retrograde flow of cadherin, thereby also promoting its retention in AJ. Reciprocal inhibition of flows would be further enhanced when tension increased the affinity of the α-catenin ABD for actin filaments, effectively increasing the mechanical load that opposed cortical flows exerted upon one another. This model assumes that regulation of the α-catenin ABD can affect the response of cortical flow to mechanical load. We tested this principle by expressing the α-catenin^A+^ mutant to increase affinity for F-actin and analysing cortical flow in cells plated on E-cadherin-coated glass, which provided a condition of uniform load from the stiff adhesive substratum. Cortical flow at the adhesive interface was decreased in cells expressing α-catenin^A+^ (Fig 5B), confirming that conformational activation of the ABD can influence cortical flow in response to mechanical load.

**Figure 5.**
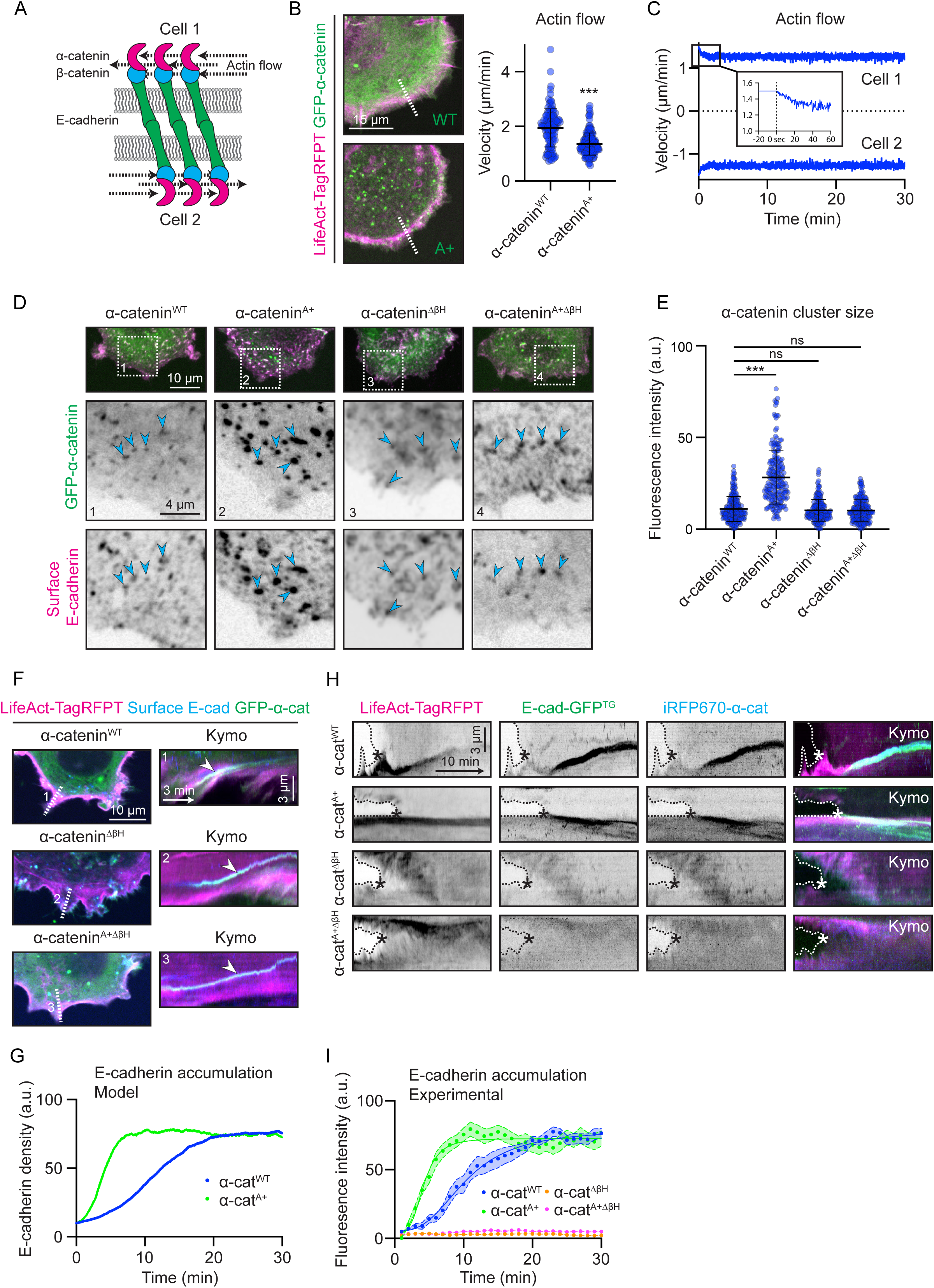
Mechanical tension activates the actin-binding domain of α-catenin to promote *cis*-clustering of E-cadherin. **(A)** Schematic representation of transligated cadherin/catenin complexes coupled to anti-parallel flowing actin cortices at cell-cell contact. **(B)** Affinity of α-catenin for F-actin influences the response of cortical flow to the mechanical load of immobilized E-cadherin. F-actin flow (LifeAct-TagRFPT, magenta) was measured in the cortical regions of MCF7 cells expressing GFP-α-catenin^WT^ or GFP-α-catenin^A+^ (green) and spread on E-cadherin ectodomain-coated substrates. Representative images and quantification of actin flow for cells expressing α-catenin^WT^ (n=120) or α-catenin^A+^ (n=142) from 3 independent experiments. **(C)** Simulations of the clutch model incorporating the effect of an α-catenin-actin bond alone on cortical flow. Top and bottom parts of graph show antiparallel velocities in each of the two cells making contact. Inset: The first seconds of the simulation, indicating a minor, transient decrease in actin flow velocity. **(D, E)** The α-catenin ABD promotes *cis*-clustering of E-cadherin. Surface unligated E-cadherin *cis*-clusters labelled with extracellular E-cadherin mAbs (magenta) in lamellipodia of non-permeabilized leader cells expressing GFP-α-catenin^WT^, GFP-α-catenin^A+^, GFP-α-catenin^ΔβH^ or GFP-α-catenin^A+ΔβH^ (green). (D) Representative images. Arrow heads: cadherin/catenin *cis-*clusters. (E) Quantification of α-catenin cluster fluorescence intensity in unligated clusters. α-catenin^WT^ clusters (n=200), α-catenin^A+^ clusters (n=200), α-catenin^ΔβH^ clusters (n=180), α-catenin^A+ΔβH^ clusters (n=200) from 4 independent experiments. **(F)** Effect of α-catenin mutants on coupling of surface E-cadherin clusters to retrograde cortical flow. Representative images and kymographs of LifeAct-TagRFPT (magenta) at the free edges of cells expressing GFP-α-catenin^WT^, GFP-α-catenin^ΔβH^ or GFP-α-catenin^A+ΔβH^ (green). Surface E-cadherin is labelled with surface-delivered E-cadherin mAbs (blue). Arrow heads: actin flow-driven retrograde movement of cadherin/catenin clusters. White dotted lines: site of kymographs. **(G)** Simulations of an enhanced clutch model that incorporates enhanced E-cadherin *cis*-clustering as an output when tension is applied to the α-catenin ABD. Model predictions for E-cadherin clutch density as a function of time. α-catenin^A+^ cells were modelled by increasing the binding rate between cadherin clutches *k_on_*, and the amount of cadherin molecules added upon α1 helix unfolding *d_add_* with respect to WT values (Table S1). **(H, I)** Enhanced cadherin *cis-*clustering by the α-catenin ABD is necessary for nascent adherens junction assembly on contact of migrating cells. E-cadherin accumulation into nascent AJ in cells expressing LifeAct-TagRFPT (magenta), E-cadherin-GFP^CRISPR^ (green) and iRFP670-α-catenin^WT^, iRFP670-α-catenin^A+^, iRFP670-α-catenin^ΔβH^ or iRFP670-α-catenin^A+ΔβH^ (blue). (H) Representative kymographs. Dotted lines: cell edges; *First moment of contact. (I) Quantification of E-cadherin accumulation (fluorescence intensity). Cell-cell contact events α-cat^WT^ (n=16), α-cat^A+^ (n=14), α-cat^ΔβH^ (n=18), α-cat^A+ΔβH^ (n=17) from 3 independent experiments. Dots: means; Light colour areas: SEM; Lines: Sigmoidal fits (α-cat^WT^, R^2^=0,67; α-cat^A+^, R^2^=0,53). ns: not significant, ***P<0,001; Mann-Whitney U test (B); Kruskal-Wallis test (E). Data are means ±SD with individual datapoints indicated.

To quantitatively evaluate this scenario, we then modelled the adherent cadherin-catenin complex as a mechanosensitive clutch where tension alters the F-actin-binding properties of the α-catenin ABD (see Computational Supplement). We define a “clutch” as a pre-coupled complex containing E-cadherin and α-catenin, *trans-*ligated to an equivalent complex on the other cell once the cells make contact. Clutches can bind to the actin cytoskeleton on both sides (Fig 5A). Upon head-on contact, initial adhesions are thus composed of a given number of clutches. Clutches get submitted to force as they bind to actin, and actin pulls in opposite directions from both cells due to the antiparallel actin flows. We then incorporated two effects of force exerted on clutches based on change in the ABD ^22, 30^. First, force can unfold the alpha1 helix, with an unfolding rate that depends on force as a slip bond. Second, unfolding of the alpha1 helix increases the affinity of the α-catenin ABD - F-actin bond. Once clutches unbind, they can rebind again according to a given binding rate.

However, our qualitative predictions were not supported in numerical simulations of the model (see Computational Supplement). Although small decreases in actin flows appeared (Fig 5C, S5B), these occurred over time scales of seconds and did not yield either the minutes-scale down-regulation of flow or adhesion growth that we observed experimentally. This was because, based on the reported K_off_ for α-catenin and F-actin ^30^, even in their high-affinity state clutches were predicted to disengage from F-actin on seconds-scales. By tuning model parameters (for instance by decreasing myosin contractility levels), the model could be adjusted to predict a substantial rather than small reduction in actin flows, but this would always occur in a time scale of seconds and not minutes. Further, the model did not predict the adhesion growth observed in experiments. This implied that enhanced binding to F-actin was not sufficient to explain how the application of tension to α-catenin could downregulate lamellipodia and promote adhesion growth.

### Tensile unfolding of the a-catenin ABD promotes the growth of E-cadherin cis-clusters and AJ assembly

We therefore considered other potential mechanisms that could be engaged when mechanical tension is applied to the ABD. In addition to increasing its affinity for F-actin, unfolding of the alpha1 helix also reveals a β-helix (βH) motif that promotes dimerization of the ABD and potentially F-actin bundling ^22^. Both these mechanisms could increase the number of cadherin-catenin complexes that are incorporated into clusters. Force-induced actin bundling is also observed in the context of cell-matrix adhesion ^31^, and its timescale (of the order of minutes rather than seconds) matches well what we observed for AJ.

To test this possibility, we asked how unfolding of the ABD influenced E-cadherin *cis*-clustering independent of other signals that are engaged by cadherin *trans*-ligation. Therefore, we measured how mutations in the α-catenin ABD affected unligated E-cadherin *cis-*clusters on the free surfaces of cells. First, we analysed the α-catenin^A+^ mutant ^22^: as this unfolds the ABD, we reasoned that it should mimic the effect of applying tension even without the mechanical loading that comes from homophilic *trans*-ligation. Indeed, expressing α-catenin^A+^ in α-catenin KO cells significantly increased the fluorescence intensity of the clusters (Fig 5D, E), consistent with enhanced recruitment of cadherin-catenin complexes. Then, we further mutated α-catenin^A+^ to delete the βH motif (α-catenin^A+ΔβH^ (S3A, B)). This abrogated the ability of α-catenin^A+^ to increase cluster size (Fig 5D, E), implying that the βH motif was necessary for α-catenin to enhance cadherin *cis*-clustering in response to tension. However, deleting the βH motif alone (α-catenin^ΔβH^ (S3A, B)) did not affect the basal levels of free clustering that were seen in cells (Fig 5D, E), although clustering was largely abolished by either α-catenin KO or mutation of the actin-binding residues within the ABD (α-catenin^A-^ (S3A, B)) (Fig S5C). Mutants lacking the βH motif were capable of binding to the cortex, as they displayed slow retrograde flow at the free surface (Fig 5F). This implied that cadherin-catenin clustering in our experiments reflected the action of two mechanisms: a basal requirement for α-catenin to bind F-actin, and an enhancement when tension unfolds the ABD to reveal the βH motif.

We then modified our clutch model so that the application of tension to cadherin clutches induced not only a higher resistance to force of α-catenin-actin bonds, but also the *cis*-growth of clusters. This was modelled by increasing the number of clutches (i.e., cadherin-catenin complexes available to bind actin) upon α-catenin ABD unfolding. Simulations of this “enhanced” clutch model now showed that cadherin junctions assembled and grew when tension was applied, with a sigmoidal profile and minutes-scale time course that closely resembled what we observed experimentally (Fig 5G, S5D). The assembly of AJ in the model was further accelerated if we increased the rate of clutch addition and the affinity of clutches to actin, thereby simulating the α-catenin^A+^ mutant (Fig 5G, S5E). This predicted that tensile activation of α-catenin could support the assembly of nascent AJ, so long as the ABD was induced to promote *cis*-clustering of E-cadherin.

These predictions of the model were supported experimentally when we measured the kinetics of nascent AJ assembly. First, nascent AJ assembly was substantially compromised when we deleted just the βH motif from otherwise-WT α-catenin (α-catenin^ΔβH^) (Fig 5H, I). This resembled the output of our original model, where AJ failed to assemble if tension did not enhance *cis*-clustering of E-cadherin. Second, AJ assembly was further increased when we reconstituted α-catenin KO cells with α-catenin^A+^, but this enhancement was lost when we deleted the βH motif (α-catenin^A+ΔβH^) (Fig 5H, I). Together, these findings show how enhanced cadherin *cis-*clustering allows tensile activation of the α-catenin ABD to mediate nascent AJ assembly.

### Enhanced cadherin-catenin cis-clustering allows mechanical antagonism to inhibit cell locomotion

Importantly, simulations also showed that cortical flow became downregulated in a sustained fashion when increased *cis-*clustering was incorporated into our enhanced model of reciprocal mechanical inhibition. Moreover, this occurred on a time scale consistent with what we observed experimentally (Fig 6A, B). We predicted that this would downregulate lamellipodial activity and allow contact to inhibit epithelial migration.

**Figure 6.**
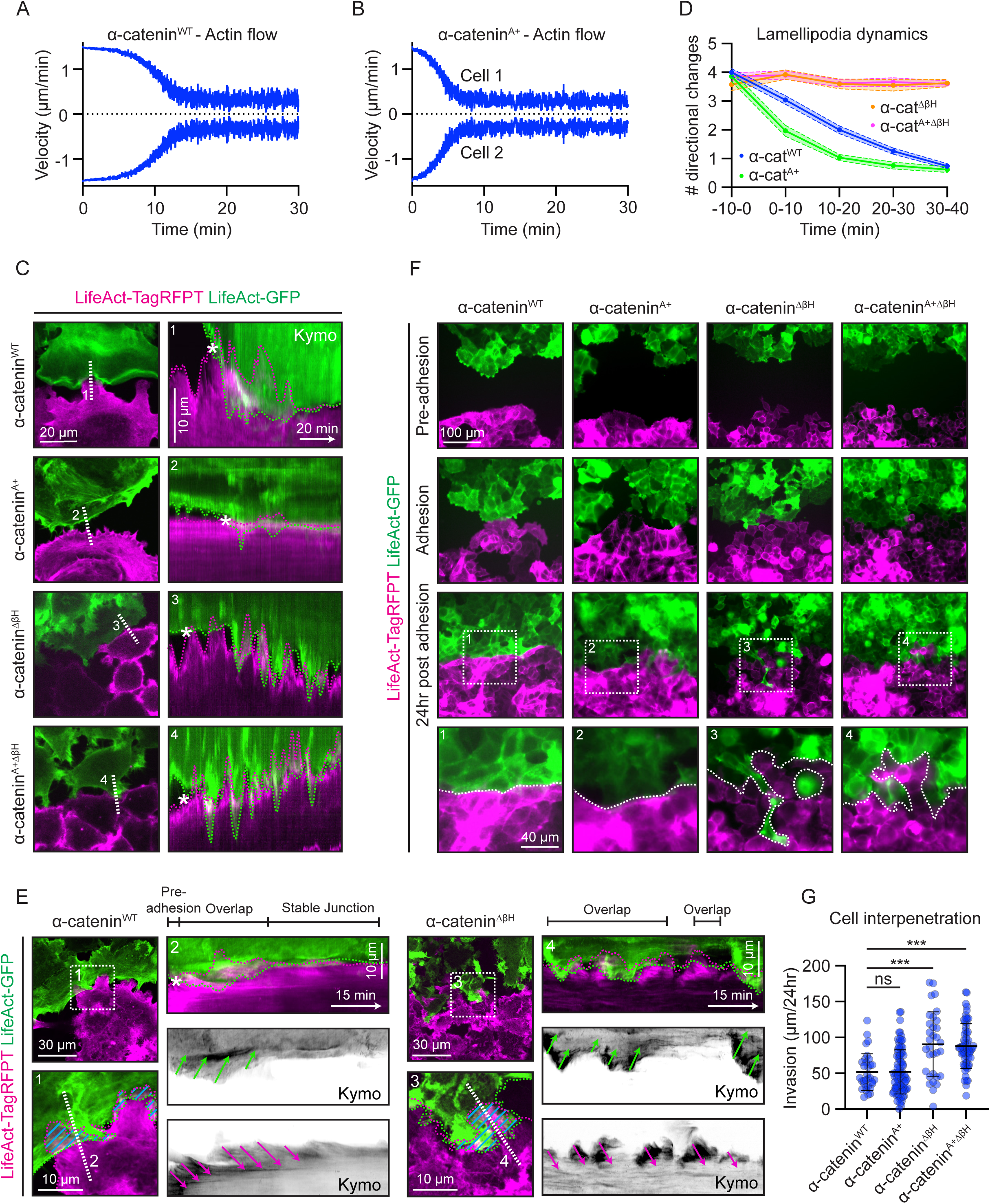
An extended bifunctional α-catenin-actin clutch allows contact to inhibit epithelial cell migration by downregulating lamellipodia. **(A)** Simulations of the clutch model for the effect of cell contact on flow velocities when tension applied by the interaction of the α-catenin^WT^ ABD with cortical flow induces *cis*-growth of E-cadherin clusters as well as increased affinity with F-actin. Top and bottom parts of graph show antiparallel velocities in each of the two cells making contact. **(B)** Simulations of the bifunctional clutch model predict that downregulation of cortical flow by contact is accelerated when the α-catenin ABD is constitutively unfolded (α-catenin^A+^). Top and bottom parts of graph show antiparallel velocities in each of the two cells making contact. **(C, D)** Cadherin *cis*-clustering by the α-catenin ABD is necessary for contact to down-regulate lamellipodial activity in expressing LifeAct-GFP (green) or LifeAct-TagRFPT (magenta). All cells express either α-catenin^WT^, α-catenin^A+^, α-catenin^ΔβH^ or α-catenin^A+ΔβH^. (C) Representative images and kymographs from movie S6. Wavy dotted lines: cell boundaries; white dotted line: site of kymograph. *First moment of contact. (D) Quantification of effect of contact on lamellipodia dynamics. Cell-cell contact events: α-cat^WT^ (n=51), α-cat ^A+^ (n=29), α-cat^ΔβH^ (n=40), α-cat^A+ΔβH^ (n=50) from 3 independent experiments. Dots: means; Light colour areas: SEM. **(E)** Cadherin *cis*-clustering by the α-catenin ABD is necessary for contact to downregulate retrograde cortical flow in cells expressing LifeAct-GFP (green) or LifeAct-TagRFPT (magenta). All cells express either α-catenin^WT^, or α-catenin^ΔβH^. Wavy dotted lines: cell boundaries; Blue lines: overlapping area; Green and magenta arrows: direction of actin flows; White dotted line: sites of kymograph. *First moment of contact. **(F, G)** Contact inhibition of locomotion measured as interpenetration of cells between WT or α-catenin KO populations expressing LifeAct-GFP (green) or LifeAct-TagRFPT (magenta). All cells express either α-catenin^WT^, α-catenin^A+^, α-catenin^ΔβH^ or α-catenin^A+ΔβH^. (F) Frames from representative movie S7. White dotted lines: cell boundaries. (G) Quantification of cell interpenetration. Cell-cell contact events: α-catenin^WT^ (n=32), α-catenin^A+^ (n=84), α-catenin^ΔβH^ (n=32), α-catenin^A+ΔβH^ (n=60) from 3 independent experiments. ns: not significant, ***P<0,001; Kruskal-Wallis test. Data are means ±SD with individual datapoints indicated.

To test this prediction, we examined how mutating the α-catenin ABD influenced the response of lamellipodia to cell-cell contact. We first reconstituted KO cells with α-catenin^A+^ to mimic the effect of an enhanced mechanical stimulus that unfolds the ABD. This accelerated the downregulation of lamellipodial activity when cells made contact with one another (Fig 6C, D; movie S6). However, it was necessary for the unfolded ABD to be capable of increasing *cis*-clustering, as the effect was lost if the βH motif was also deleted (α-catenin^A+ΔβH^) (Fig 6C, D; movie S6). Furthermore, deleting the βH motif alone (α-catenin^ΔβH^) abolished the capacity for cell-cell contact to inhibit lamellipodia (Fig 6C, D; movie S6). Consistent with our model of reciprocal inhibition, the ability of contact to inhibit retrograde flow was also disrupted when we deleted the βH motif (Fig 6E). Therefore, enhanced *cis*-clustering allows tensile activation of the α-catenin ABD to inhibit cortical flow and downregulate lamellipodial activity.

Finally, we asked whether the molecular effects of the βH motif translated to affect contact inhibition of epithelial migration. We found that colliding α-catenin^ΔβH^ populations failed to inhibit cell migration. As revealed by labelling different populations with differently-tagged LifeAct, α-catenin^ΔβH^ populations continued to interpenetrate one another upon contact, rather than forming the relatively smoothly defined borders seen in WT populations (Fig 6F, G; movie S7). The α-catenin^A+^ mutant did not increase the robust degree of contact inhibition already displayed by the WT cells; however, its ability to mediate contact inhibition was also disrupted by deleting the βH motif (α-catenin^A+ΔβH^) (Fig 6F, G; movie S7). Altogether, these findings indicate that tensile activation of α-catenin ABD allows cell-cell contact to inhibit epithelial migration by downregulating lamellipodia.

## DISCUSSION

In this study, we sought to understand how migrating epithelial cells recognize when to stop moving when they come into contact with one another. Specifically, we endeavoured to identify the proximate mechanism by which E-cadherin adhesion inhibits epithelial locomotion. We now propose the following model (Fig 7A): The trigger for CIL occurs when E-cadherin complexes on the free surfaces of migrating cells undergo *trans*-ligation and mechanically connect together the anti-parallel, retrograde cortical flows of colliding lamellipodia. This applies a tensile signal that engages the α-catenin ABD to recruit further cadherin-catenin complexes into adhesions as well as increases its affinity for actin filaments. Adhesion growth further enhances the mechanical coupling between the antiparallel cortical flows, leading to their reciprocal downregulation. Loss of cortical flow then compromises the lamellipodial activity of leader cells to inhibit epithelial migration. Thus, in this model mechanosensing by the α-catenin ABD allows cells to interpret the topology of cortical flows, as a basis for identifying which contacts will inhibit locomotion.

**Figure 7.**
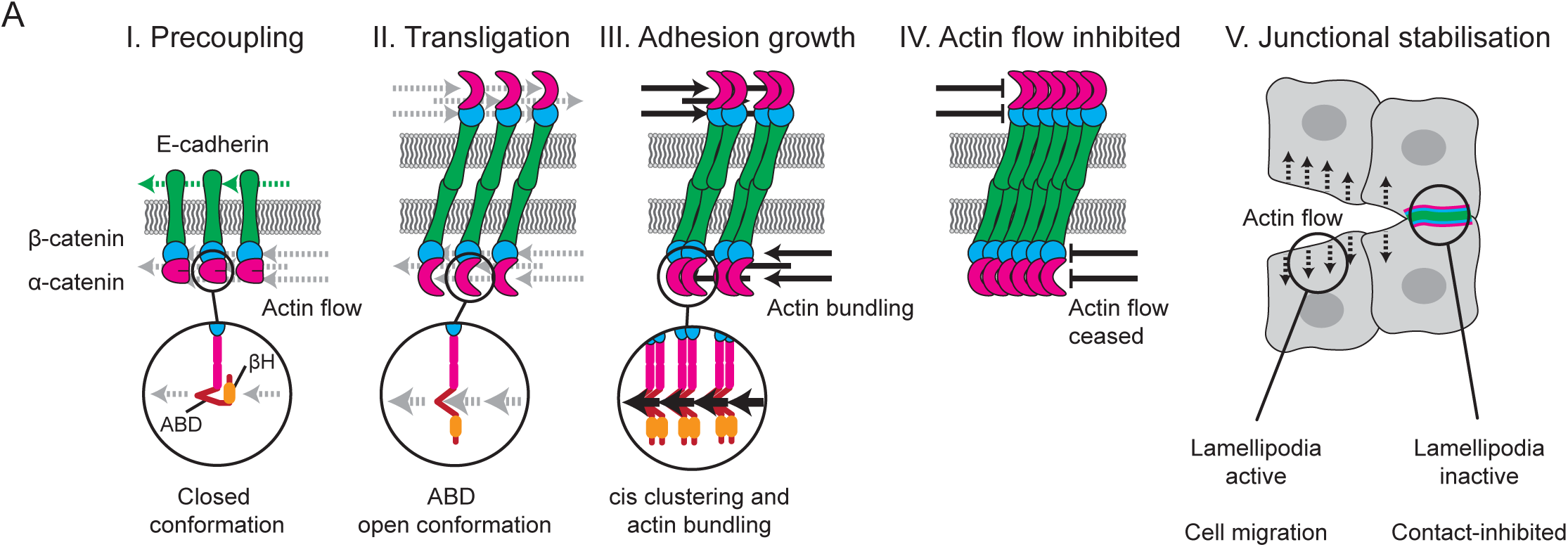
α-catenin clutch-induced AJ assembly inhibits lamellipodial activity and cell migration. **(A)** Schematic overview of how the α-catenin clutch induces AJ assembly. I) Surface E-cadherin clusters are coupled to actin retrograde flow through β- and α-catenin. II) E-cadherin trans-ligation results in significant increase in load caused by anti-parallel actin flows. As a result, the α-catenin actin binding domain (ABD) changes to an open conformation. III) This conformational change increases the α-catenin binding affinity for actin and causes cis-clustering through the α-catenin βH-domain, resulting in the bundling of actin filaments and growth of the newly formed AJ. IV) Cis-clustering of cadherin-catenin complexes inhibits cortical actin flow. V) As a result, the formation of lamellipodia ceases and further cell migration is blocked.

A role for E-cadherin in mediating contact inhibition of epithelial migration was suggested by the observation that the locomotor activity of lamellipodia decreased as nascent AJ assembled. This was supported by the finding that α-catenin KO prevented contact from inhibiting either epithelial migration as a whole or the lamellipodial activity of the colliding leader cells. Several lines of evidence argue that tensile activation of the α-catenin ABD plays a key role in allowing cadherin adhesion to inhibit locomotion. First, mechanical tension across α-catenin was greatest at the head-on contacts that inhibit epithelial migration, as revealed by antibodies that detect tension-sensitive conformational change in α-catenin. Second, several scaffolding domains in the α-catenin molecule respond to increased mechanical tension ^22, 25, 30, 32^. However, mimicking tensile unfolding of the ABD with the α-catenin^A+^ mutant was sufficient to enhance the ability of head-on contacts to coordinately promote AJ assembly while inhibiting retrograde flow and lamellipodial activity. Third, CIL required the ability of the ABD to promote *cis-*clustering of cadherin complexes, via dimerization of α-catenin molecules and potentially also bundling of actin filaments. This effect was first identified for the βH motif of the ABD based on *in vitro* studies ^22^; it was confirmed in our work by the observation that this motif allowed the unfolded ABD to enhance *cis-*clustering of cadherin-catenin complexes even on the free surfaces of cells. Importantly, the ability of contact to inhibit migration and lamellipodial activity was abolished when the βH motif of the ABD was deleted in either WT α-catenin or the tension-mimicking α-catenin^A+^ mutant.

Thus, although increased affinity for actin filaments is the best-understood consequence of applying tension to the α-catenin ABD ^22, 30, 33^, *cis-*clustering of cadherin-catenin complexes is a second effect that is essential for contact to inhibit epithelial locomotion. The α-catenin ABD can then be considered as a bi-functional tension-activated scaffold, which supports distinct mechanisms that increase affinity to F-actin and enhance the recruitment of cadherin complexes into nascent adhesions. Of note, although deletion of the βH motif was reported to affect F-actin binding *in vitro* ^22^, this was not apparent at the cellular level, as tested by measuring the coupling of free cadherins to cortical flow. Our results do not exclude the possibility that other α-catenin partners contribute at later stages of contact inhibition. Nonetheless, they argue strongly that activation of the ABD represents the first, key stage in the process.

Ultimately, cadherin adhesions must inhibit epithelial migration by acting on some critical element in the locomotor apparatus of the cells. Retrograde flow is an important driver of cell locomotility, which generates shear forces for cell movement ^17, 18^. Therefore, it was noteworthy that the inhibition of lamellipodial activity at contacts coincided with downregulation of retrograde cortical flow. Changes in retrograde flow also exactly paralleled how manipulating the α-catenin ABD affected contact inhibition of either lamellipodia or epithelial migration. Thus, contact inhibition of retrograde flow was accelerated by expression of the constitutively-opened α-catenin^A+^ mutant, while contact failed to inhibit retrograde flow when the βH motif was deleted. This is consistent with the notion that α-catenin in the cadherin complex allowed cell contact to inhibit lamellipodia by downregulating retrograde cortical flow. Along these same lines, disrupting the ability of contact to inhibit retrograde flow by deleting the βH motif caused colliding cell populations to interpenetrate, evidence that CIL required that cadherin adhesion downregulate cortical flow. Together, these findings identify control of cortical flow in lamellipodia as key for cadherin adhesions to inhibit epithelial migration.

How could activation of the α-catenin ABD allow cadherin adhesion to downregulate cortical flow? One parsimonious explanation is where antiparallel cortical flows reciprocally inhibit one another by mechanical loading. The capacity for enhanced mechanical loading to inhibit flow was demonstrated in principle by finding that expression of α-catenin^A+^ downregulated the cortical flows that cells made at adhesions with cadherin-coated substrata. It was reinforced by the observation that flows were not inhibited at heterologous contacts between WT and α-catenin KO cells, a situation where antiparallel flows could not act upon one another because one side of the *trans*-ligated cadherin-catenin complex was unable to bind F-actin. Note that in this model, *trans-*ligated cadherin complexes principally serve as mechanical couplers of the flows; and conformational activation of the α-catenin ABD would exert its effects by increasing mechanical coupling, thereby increasing the load upon flows. Although increasing the affinity of α-catenin ABD for actin filaments is expected to facilitate mechanical coupling, our computational modelling suggested that this mechanism alone was not sufficient to inhibit cortical flow on the time scales that we observed experimentally. Instead, sustained inhibition of flow required that the tension-activated ABD also promote adhesion growth, as confirmed by deleting the βH motif.

There is an attractive simplicity to this mechanical model. It has the capacity to exert rapid effects: since cadherins already interacted with the cortex on the free surfaces of cells, they were poised to couple flows upon contact. Furthermore, this model has the potential for subcellular spatial specificity, as flows will only be inhibited at cadherin adhesions that are exposed to sufficient tension. This helps to explain why epithelia can migrate as coherent sheets coupled together by E-cadherin adhesion, yet inhibit migration when they collide head-on and generate greater tensional signals. These design features would allow mechanical inhibition to serve as the trigger for contact to inhibit epithelial migration. Our mechanical model does not, however, exclude contributions from other tension-sensitive signaling events. For example, mature AJ in MCF7 and other epithelial cells are characterized by a contractile cortex ^27, 34^. Although Myosin II was not recruited into nascent AJ until after the initial phase of lamellipodial inhibition (not shown), it is possible that the subsequent assembly of a Myosin-enriched cortex replaces the tensile forces of retrograde flow to sustain contact inhibition of locomotion. As well, cadherin adhesion has complex effects on Arp2/3, the principal actin nucleator at the leading edges of lamellipodia. VE-cadherin junctions are reported to downregulate Arp2/3-based actin assembly ^35^, whereas Arp2/3 can also be recruited to E-cadherin to nucleate actin at AJ ^36^. Whether tensional signals generated by coupling antiparallel flows can also regulate Arp2/3 is an interesting question for future research.

In conclusion, we propose that the bifunctional clutch of the α-catenin ABD allows the cadherin system to interpret tensional stimuli of cortical flows to inhibit epithelial cell locomotion. This emphasizes the key role that topology of cortical flow plays in our model. First, the tug-of-war topology of flow that is found at head-on contacts can explain why tension across the cadherin complex is greater at this site than at side-side contacts, where flows have a parallel orientation across the junction. This tensile difference causes the α-catenin ABD clutch to be activated preferentially at head-on, compared to side-side contacts. Second, reciprocal inhibition of lamellipodia occurs when the mechanical coupling of anti-parallel flows is further enhanced as the activated clutch promotes adhesion growth. In other words, the anti-parallel topology of flows is initially responsible for generating a tensional signal strong enough to activate the α-catenin ABD and later mediates reciprocal inhibition when the flows are coupled by cadherin-catenin clutches. This model does not exclude contributions for other contact-based cell surface signals to CIL ^37^. However, it has ideally placed cells to make early-immediate responses to contact by coupling the molecular apparatus of adhesion to the cytoskeletal driver of lamellipodia. Since patterns of cortical flow reflect the overall polarity of migrating cells, this mechanosensitive apparatus allows migrating epithelia to evaluate the orientation of cells with which they collide, discriminating head-on from side-to-side contacts, and alter locomotion accordingly.

### Caveats

We used the MCF7 system for our experiments, a well-characterized model of epithelial migration where AJ inhibit locomotility when they assemble at head-on collisions. This system also reproduces many features of locomoting epithelia, including dynamic lamellipodia and a requirement for AJ to also mediate sheet cohesion. It will now be important to test how the model that we have identified may operate in other epithelia, notably in vivo models of sheet migration.

## Supporting information

Key recources table

## ACKNOWLEDGEMENTS

We thank Srikanth Budnar for the E-cadherin-GFP^CRISPR^ cell line, Ivanka Herat and Suzie Verma for technical assistance, and all our colleagues in the lab for their support, advice and encouragement. The work in Australia was supported by grants (GNT1163462, 2010704 to A.S.Y.; GNT1158002 to E.G) and fellowships (GNT1136592) from the National Health and Medical Research Council of Australia and the Australian Research Council (DP19010287, 190102230 to AY and DP200100737 to E.G). E.G. was supported by a Future Leader Fellowship – Level 2 (104692) from the National Heart Foundation of Australia; RGM by EMBL Australia; I.N. by the European Molecular Biology Organization (EMBO ALTF 251-2018); and L.S. by a UQ Early Career Researcher Grant (UQECR2058733). Microscopy was performed at the ACRF/IMB Cancer Research Imaging Facility created with the generous support of the Australian Cancer Research Foundation.

P.R.C. and M.D.H. acknowledge funding from the Spanish Ministry of Science and Innovation (PID2019-110298GB-I00), the European Commission (H2020-FETPROACT-01-2016-731957) the Generalitat de Catalunya (2017-SGR-1602), The prize ‘ICREA Academia’ for excellence in research to P.R.C., Fundació la Marató de TV3 (201936-30-31 to P.R.C.) and ‘la Caixa’ Foundation (grant LCF/PR/HR20/52400004). The Institute for Bioengineering of Catalonia (IBEC) is a recipient of a Severo Ochoa Award of Excellence from MINCIN. D.V. acknowledges funding by the Deutsche Forschungsgemeinschaft (CRC1348, B01). S.M. acknowledges the support of Department of Atomic Energy, Government of India, under Project Identification No. RTI 4006 and NCBS-TIFR-core funds, the J.C. Bose Fellowship from DST, Government of India, and India Alliance DBT - Wellcome Trust Margdarshi fellowship (IN M/15/1/502018).

## AUTHOR CONTRIBUTIONS

Conceptualization, I.N., P.R.C., A.S.Y.; Methodology, I.N., M.D.H., P.R.C.; Formal analysis, I.N., M.D.H.; Investigation, I.N., L.S., A.B-M., D.C-R., C.N.D; Writing, I.N., M.D.H., J.M.K., R.G.M., D.V., S.M., E.G., P.R.-C., A.S.Y.; Visualization, I.N., M.D.H., D.C-R; Supervision, R.G.M., P.R.-C., A.S.Y.; Funding acquisition, I.N., L.S., D.V., S.M., E.G., P.R.-C., A.S.Y.

## STAR METHODS

### RESOURCE AVAILABILITY

#### Lead contact

Further information and requests for resources and reagents should be directed to and will be fulfilled by the lead contacts, Alpha Yap: (experimental) and Pere Roca Cusachs: (theory).

#### Materials availability

All reagents generated in this study will be made available upon request to the lead contact.

#### Data and Code Availability

All data and code generated in this study will be made available upon request to the lead contact.

### EXPERIMENTAL MODEL AND SUBJECT DETAILS

#### Cell culture and transfection

MCF7 cells were obtained from ATCC (HTB-22) and cultured in Dulbecco’s Modified Eagle Medium (DMEM) supplemented with 10% fetal bovine serum (FBS) (GE Healthcare Australia), 100 units/ml penicillin/streptomycin (pen/strep) (Life Technologies Australia) and 1 mM CaCl2. HEK293T cells were cultured in DMEM with L-glutamine and sodium pyruvate (Invitrogen), supplemented with 10% FBS and 100 units/ml Pen/Strep. All cells were cultured at 37°C in 5% CO2.

Lipofectamin 3000 (Thermo Fisher) was used to transfect MCF7 cells for ectopic expression of transgenes and CRISPR gene-editing constructs. PEI 2500 (BioScientific) was used to transfect HEK293T cells for lentiviral packaging. Both methods were performed according to manufacturer’s protocols.

#### Stable cell lines

MCF7 E-cadherin-GFP knock-in cells were generated using CRISPR-Cas9 technology and were described before ^38^. MCF7 α-catenin knockout cells were generated using CRISPR-Cas9 technology. Cells were transfected with the pSpCas9(BB)-2A-Puro (PX459) vector bearing the appropriated sequence (5’-GAAATGACTGCTGTCCATGC-‘3) to target *CTNNA*1 (α-catenin). 24 hours after transfection, cells were subjected to 0,8 µg/ml puromycin for 48 hours. After selection, single clones were grown and α-catenin expression was assessed using western blot and IF staining. Genomic sequencing was performed to confirm knockout.

MCF7 cells expressing LifeAct-GFP, LifeAct-TagRFPT, LifeAct-iRFP670, E-cadherin-GFP^TG^, iRFP670-α-catenin^WT^, iRFP670-α-catenin^A+^, iRFP670-α-catenin^ΔβH^ and iRFP670-α-catenin^A+ΔβH^ were generated using lentiviral transduction. Lentiviral constructs were packaged into lentivirus in HEK293T cells by means of third generation lentiviral packaging plasmids. Lentivirus containing supernatant was harvested on days 2 and 3 after transfection, concentrated using a Lenti-X concentrator (Clontech #631232) and used to transduce target cells. Lentiviral-transduced MCF7 cells were subjected to either 0,5 µg/ml puromycin or 400 µg/ml G418 for 72 hours. Subsequently, cells were maintained in either 0,25 µg/ml puromycin or 50 µg/ml G418.

### METHOD DETAILS

#### DNA constructs

pLVX-IRES-Puro (PLViP) and pLVX-IRES-Neo (PLViN) were used for lentiviral packaging. PLViN was constructed by replacing the EF1a promotor for a CMV promotor in pLV-EF1a-IRES-Neo (Addgene #85139). PLViP-LifeAct-GFP and PLViP-LifeAct-TagRFPT were constructed by replacing iRFP670 for GFP- and TagRFPT-coding sequences in PLViP-LifeAct-iRFP670 (Kindly provide by Dr. J. van Buul, Sanquin Research and Landsteiner Laboratory, The Netherlands). pLViN-E-cadherin-GFP^TG^ was constructed by replacing YFP for a GFP-coding sequence in N1-E-cadherin-YFP. Subsequently, E-cadherin-GFP was cloned into PLViN. GFP-α-catenin^WT^ was constructed by cloning a GFP-coding sequence into acat 1-906 (Addgene #24194). GFP-α-catenin^A+^ (RAIM670-733GSGS), GFP-α-catenin^A-^ (L785A I792A V796A) and GFP-α-catenin^V-^ (L344P) were constructed by Gene Universal using GFP-α-catenin^WT^. GFP-α-catenin^ΔβH^ (ΔE795-810S) was constructed using GFP-α-catenin^WT^. GFP-α-catenin^A+ΔβH^ (RAIM670-733GSGS, ΔE795-810S) was constructed using GFP-α-catenin^A+^. All α-catenin mutants were properly incorporated into adherens junctions after expression in WT MCF7 cells, indicating β-catenin binding was not affected (Fig S3C). PLViN-iRFP670-α-catenin^WT^, PLViN-iRFP670-α-catenin^A+^, PLViN-iRFP670-α-catenin^ΔβH^ and PLViN-iRFP670-α-catenin ^A+ΔβH^ were constructed by replacing GFP for an iRFP670-coding sequence in the GFP-α-catenin constructs. Subsequently, iRFP670-α-catenin constructs were cloned into PLViN. All constructs were confirmed by sequencing.

#### Antibodies, drugs and chemicals

Mouse monoclonal antibodies against E-cadherin ectodomain (#563571) (used for IF) and β-catenin (#610154) were purchased from BD Biosciences. Mouse monoclonal antibody against E-cadherin (used for Western) was kindly provided by Dr. M. Takeichi (Riken, Japan). Rabbit polyclonal antibody against GAPDH (#2275-PC-100) was purchased from R&D Systems. Rabbit polyclonal antibody against α-catenin (#71-1200) (used for Western blot) was purchased from Thermo Fisher. Rabbit polyclonal antibody against α-catenin (used for IF) was kindly provided by Dr. B. M. Gumbiner (Seattle Children’s, USA). Rat monoclonal antibodies against E-cadherin were purchased from Abcam (#ab11512) (used for IF) and Thermo Fisher (#13-1900) (used for IF). Rat monoclonal antibody against α-catenin M-domain (α18) was kindly provided by Dr. A. Nagafuchi (Kumamoto University, Japan). Rabbit polyclonal antibody against a-catenin ABD (VD7) has been described before ^26^.

Alexa-Fluor-405- (#A-31553, #A-48254, #A-48261), Alexa Fluor-488- (#A-11001, #A-11008, #A-11006), Alexa-Flour-594- (#A-11032, #A-11037, #A-48264) and Alexa Fluor-647- (#A-21236, #A-21245, #A-48265) conjugated goat antibodies against mouse, rabbit and rat were purchased from Thermo Fisher. For Western blot, goat anti-mouse-HRP conjugate (#1706516) and goat anti-rabbit-HRP conjugate (#1706515) were purchased from Bio-Rad.

Alexa Fluor® 488- (A12379), 594- (A12381) and 647- (A22287) conjugated phalloidin were purchased from Thermo Fisher. SiR-Tubulin (CY-SC002) was purchased from Cytoskeleton and used at a concentration of 1 µM. Cells were incubated with SiR-Tubulin for 1 hour prior to imaging. Para-nitroblebbistatin (DR-N-111) was purchased from Optopharma and used at a concentration of 10 µM. CK666 (SML0006) was purchased from Sigma-Aldrich and used at a concentration of 50 µM. Jasplakinolide (420107) was purchased from Merck and used at a concentration of 100 nM. In order to stop cortical actin flow, cells were treated with Para-nitroblebbistatin for 1 hour, following by 2-hour treatment with Para-nitroblebbistatin, CK666 and Jasplakinolide (BCJ).

#### E-cadherin-coated substrate assay

Coverslips were covered with 25 µg/ml Protein A (Thermo Fisher, #21181) and incubated at room temperature (RT) for 3 hours. After this step, the coverslips were kept wet. Coverslips were washed with PBS and 25 µg/ml hE-cadherin/Fc (SinoBiological, #10204) was added, followed by overnight (ON) incubation at 4°C. Following incubation, coverslips were washed with PBS and incubated with 1% BSA for 45 minutes at RT. Finally, the prepared substrates were washed with PBS and movie media (15 mM HEPES, 5 mM CaCl_2_, 2 mM L-glutamine and 0,05% FCS) was added.

Soft PDMS substrates with a stiffness of 2-3 kPa were prepared by mixing CyA and CyB (Dow Corning) at a 1:1 ratio. The substrates were cured at 80°C for 2 hours before silanizing with 5% 3-aminopropyl trimethoxysilane (Sigma Aldrich, #281778) in ethanol. 0,05% of 200nm carboxylated fluorescent beads (Thermo Fisher, #F8807) were added to the substrates and the substrates were coated with protein A and hE-cadherin/Fc as described above.

To perform E-cadherin-coated substrate spreading assays, cells were trypsinized with Crystalline Trypsin (Sigma Aldrich, #T-0303) just long enough to see the first cells rounding up (6 minutes for MCF7 cells). Cells were resuspended in movie media and centrifuged for 3 minutes at 0,5 G. Next, supernatant was removed, cells were resuspended and filtered through a 40 µm cell strainer to generate single cells. Lastly, cells were added to the prepared E-cadherin-coated substrates and spreading was monitored for 2 hours before additional treatments were performed.

#### Trypsin protection assay

Cells were consecutively washed with Hanks’ Balanced Salt Solution (HBSS) and incubation media (WCL, C+: 2 mM CaCl_2_ in HBSS; C-: 2 mM EGTA in HBSS) without trypsin. Next, incubation media with trypsin was added to the cells (WCL: 2 mM CaCl_2_ in HBSS; C+: 2 mM CaCl_2_ + 0,5% Crystalline Trypsin in HBSS; C-: 2 mM EGTA + 0,5% Crystalline Trypsin in HBSS) and cells were incubated for 20 minutes at 37°C. Media and floating cells were transferred to tubes and 1 ml cold (4°C) 2mM CaCl_2_ + 1% FCS in HBSS was added to the cells. Cells were scraped and transferred to the same tubes, followed by maximum speed centrifugation for 2 minutes at 4°C. Finally, supernatant was removed and sample buffer + DTT was added to the cells to prepare them for Western blot analysis.

#### Western blotting

Cell extracts were prepared in sample buffer (62,5 mM Tris-HCl pH6,8; 2,5% SDS; 0,002% Bromophenol Blue; 5% DTT, 8% glycerol), boiled for 5 minutes, and loaded on SDS-PAGE gels. After running the gels, samples were transferred onto 0,22 µm-pore membranes. Next, blots were blocked with 5% BSA, incubated ON with primary antibodies at 4°C and incubated for 1 hour with HRP-conjugated secondary antibodies at RT. After incubation, blots were treated with Supersignal West Pico PLUS Chemiluminescent Substrate (Thermo Fisher, #34579) and imaged on a Bio-Rad ChemiDoc MP.

#### Immunofluorescence staining

Cells were fixed with either −20°C MeOH for 10 minutes (E-cadherin, β-catenin, α-catenin, α18, VD7) or 4% PFA for 20 minutes at RT (E-cadherin, phalloidin), followed by permeabilization with 0,2% Triton X-100 for 1 minute. Next, samples were blocked with 1% BSA dissolved in PBS supplemented with 0,05% Tween-20 for 45 minutes and sequentially incubated with primary antibodies for 1 hour and fluorescently labelled secondary antibodes and/or phalloidins for 45 minutes. Finally, samples were washed, dried and mounted in ProLong Gold with (Cell Signaling, #8961) or without (Cell Signaling, #9071) DAPI.

#### Microscopy

Fixed and live samples (FRAP) were imaged on a Leica DMi8 inverted confocal microscope equipped with 405nm, 440nm pulsed, 442nm and 470nm – 670nm white light lasers and HyD detectors. Images were acquired using a 100x (1,40 N.A.) oil HC Plan Apochromat objective.

Fixed and live samples were imaged on an Andor Dragonfly Spinning Disc inverted confocal microscope equipped with 405nm, 445nm, 488nm, 514nm and 637nm lasers and Zyla 4.2 sCMOS detectors. Images were acquired using 40x (0,95 N.A.) air or 60x (1,4 N.A.) oil Plan Apo λ objectives.

Live samples were imaged on a Zeiss AxioObserver upright widefield microscope equipped with LED illumination and sCMOS detector. Images were acquired using a 20x (0,5 N.A.) air EC Plan-Neofluar objective.

Fixed samples were imaged on a Zeiss AxioImager upright widefield microscope equipped with LED illumination and CCD detector. Images were acquired using a 63x (1,4 N.A.) oil Plan Apochromat objective.

#### Image analysis

Lamellipodia dynamics was analysed using movies of cells expressing LifeAct-GFP and LifeAct-TagRFPT. Directional changes of the lamellipodium edge were quantified (Fig S1B) and grouped in 10 minute bins. T=0 indicates the first frame in which lamellipodia made contact. Kymographs were generated using the ‘KymoRescliceWide’ ImageJ plugin with a linewidth of 1 µm.

E-cadherin accumulation was analysed using movies of cells expressing LifeAct-TagRFPT and E-cadherin-GFP^CRISPR^. E-cadherin-GFP^CRISPR^ fluorescence intensity was measured in 3,17 µm^2^ circular regions of interest in which the earliest adhesions appeared. T=0 indicates the first frame in which lamellipodia made contact, based on the LifeAct-TagRFPT signal. Background was subtracted and all values were normalized before graphs were plotted. Sigmoidal functions were fitted to data points. Kymographs were generated using the ‘KymoRescliceWide’ ImageJ plugin with a linewidth of 1 µm.

Cell invasion was analysed using movies of cells expressing LifeAct-GFP and LifeAct-TagRFPT. Invasion was measured 24 hours after first contact in 166 µm wide regions of interest.

Relative numbers of intracellular and surface E-cadherin clusters were analysed in images of cells labelled for total and surface E-cadherin using IF. Gaussian blur was applied to images and clusters were detected using automated ‘Moments’ thresholding followed by particle detection using particle analysis without a size cut-off. Intracellular cluster pool was calculated by subtracting the surface cluster pool from the total cluster pool.

Cadherin/catenin cluster velocities were analysed using movies of cells expressing E-cadherin-GFP^CRISPR^, E-cadherin-GFP^TG^, GFP-α-catenin, iRFP670-α-catenin or by E-cadherin ectodomain-binding antibodies to label E-cadherin surface clusters. Only clusters that could be tracked from start to end within the time of recording were quantified. In the case of a gradual decrease of velocity, average velocity per cluster was defined by drawing a straight line from start to end in the corresponding kymograph. Actin flow velocities were analysed using movies of cells expressing LifeAct-GFP, LifeAct-TagRFPT or LifeAct-iRFP670. Kymographs were generated using the ‘KymoRescliceWide’ ImageJ plugin with a linewidth of 0,7 µm.

α-catenin conformational changes were visualized by α18/α-catenin IF labelling (to analyse the M-domain), or VD7/α-catenin IF labelling (to analyse the ABD). Ratiometric images were generated by dividing the α18 or VD7 images by the total α-catenin images. Gaussian blur was applied to images before calculation. For quantification, the ratiometric signal was measured at the junctions of interest. For anti-parallel flow junctions at head-on contacts, cells were fixed between 1 and 2 hours after first contact.

FRAP measurements were performed by bleaching 8-µm diameter circles in the region of interest, followed by 6:40 minutes of imaging with a frame interval of 2 seconds. Logistic functions were fitted to data points

α-catenin cluster sizes were analysed in images of cells expressing GFP-α-catenin mutants and labelled for surface E-cadherin using IF. GFP-α-catenin fluorescence intensity was measured in 0,82 µm^2^ circular regions of interest in which an E-cadherin surface cluster was visible. Background was subtracted to correct for expression levels.

#### Software

All images and movies were analysed, and all movies were labelled using ImageJ 1.52. All plots were generated, and all statistical tests were performed in Prism 9.0.0. All modelling was performed using Matlab 9.9 (R2020b). The Leica DMi8 inverted confocal microscope was controlled by LASX software. The Andor Dragonfly Spinning Disc inverted confocal microscope was controlled by Fusion software. The Zeiss AxioImager upright widefield microscope and the Zeiss AxioImager upright widefield microscope were controlled by Zen Blue 3 software.

### QUANTIFICATION AND STATISTICAL ANALYSIS

Statistical parameters for individual experiments are listed in the corresponding figure legends. This includes the statistical test performed, sample size (N), number of independent experiments and statistical significance.

## SUPPLEMENTAL FIGURE LEGENDS

**Figure S1.**

**(A)** Junction formation between MCF7 cells expressing LifeAct-TagRFPT (magenta) and E-cadherin-GFP^CRISPR^ (green). White dotted line: free cell edge; arrow heads: Adherens Junctions (AJ).

**(B)** Analysis example lamellipodia dynamics. Every directional change in the kymograph (switch from protrusive to retractive motion or vice versa) is counted as 1.

**(C)** Genomic sequencing results of CTNNA1, exon 1, in α-catenin KO cells.

**(D)** Western blot analysis of MCF7 WT and α-catenin KO cells.

**(E)** Confluent layer of WT and α-catenin KO cells stained for α-catenin (green) and E-cadherin (magenta).

**(F)** Trypsin protection assay and western blot analysis of WT and α-catenin KO cells. Surface E-cadherin is protected against trypsin cleavage by the addition of Cacl_2_. WCL = Whole Cell Lysate, C+ = with Cacl_2_ (Total E-cadherin levels), C- = without Cacl_2_ (Intracellular E-cadherin only).

**Figure S2.**

**(A)** MCF7 leader cells expressing LifeAct-TagRFPT (magenta) and E-cadherin-GFPCRISPR (green). Wavy white dotted line: free cell edge; straight white dotted line: site of kymograph; white arrow heads: retrograde movement (Kymo) of E-cadherin clusters.

**(B)** Leader cells expressing E-cadherin-GFP^TG^ (green), LifeAct-TagRFPT (magenta), stained for microtubules using SiR-Tubulin (blue in overview, magenta in zooms). White arrow heads: E-cadherin vesicle moving over a microtubule.

**(C)** Leader cells expressing LifeAct-TagRFPT (magenta) and E-cadherin-GFP^TG^ (green). White and blue arrow heads: E-cadherin cluster formation in the lamellipodia.

**(D)** Leader cells stained for E-cadherin (green), β-catenin (magenta) and α-catenin (blue).

**(E)** Leader cells expressing LifeAct-TagRFPT (magenta) and E-cadherin-GFP^TG^ (green). Images show effect of BCJ treatment on E-cadherin and actin. White dotted line: site of kymograph (2E).

**(F, G)** WT and α-catenin KO leader cells stained for total E-cadherin (green) and surface E-cadherin (magenta). (F) Representative images. Blue arrow heads: E-cadherin surface clusters. (G) Quantification of E-cadherin surface clusters. WT cells (n=18), α-catenin KO cells (n=20) from 2 independent experiments.

***P<0,001; Mann-Whitney U test. Data are means ±SD with individual datapoints indicated.

**Figure S3.**

**(A)** Domain organization, mutation indications and binding efficiencies of α-catenin^WT^ and all α-catenin mutants used in this study.

**(B)** Localization of GFP-α-catenin transgenes (green) in WT cells co-stained for E-cadherin (magenta) and actin (blue). All α-catenin mutants are properly incorporated into adherens junctions, indicating β-catenin binding is not affected.

**Figure S4.**

**(A)** Schematic representation of a cell spread on an E-cadherin ectodomain-coated PDMS substrate and the effect of actin flow on the cadherin-catenin complexes.

**Figure S5.**

**(A)** MCF7 α-catenin KO cells expressing GFP, GFP-α-catenin^WT^, GFP-α-catenin^A+^, GFP-α-catenin^A-^, GFP-α-catenin^V-^, GFP-α-catenin^ΔβH^ or GFP-α-catenin^A+ΔβH^ (green), stained for E-cadherin (magenta) and actin (blue). White dotted line: free cell edge.

**(B)** Force evolution corresponding to (5C)

**(C)** Surface E-cadherin clusters in leader WT cell, α-catenin KO cell and α-catenin KO cell reconstituted with GFP-α-catenin^A-^ (green).

**(D)** Force evolution corresponding to (6A)

**(E)** Force evolution corresponding to (6B)

**Figure S6.**

**(A)** Scheme of the molecular ensemble. Actin filaments of neighboring cells are connected via a set of CCC-CCC links. CCC from one cell bind through cadherin trans interaction to the equivalent CCC from the other cell. α-catenin-F-actin bond regulates force transmission between actin filaments throughout the whole link.

**(B)** Dynamics of the clutch. As regards F-actin – α-catenin bond, it can either unbind with a *k_off_*(*F*), bind according to *k_on_*(*d_CCC_*) or unfold following *k_unf_*(*F*).

**(C)** Time evolution of the system. I Initial random distribution of clutches, some engaged and others not. II As time passes, actin filaments begin to move in opposite directions, thereby loading some force on the system due to the tension induced on the springs. III As the actin filaments continue to pull, deformation on the spring becomes greater, and so it does the total loaded force, decreasing the actin filament movement. IV High tension on the links induces an increase on the number of clutches in the system, thereby generating a loaded force able to completely stop the flow.

## SUPPLEMENTAL MOVIE LEGENDS

**Movie S1.** Lamellipodia dynamics and actin flows during contact formation of MCF7 WT cells **(A)**, **related to Figure 1C and 3B**, or α-catenin KO cells **(B), related to Figure 1G and 3D**, expressing LifeAct-GFP (green) or LifeAct-TagRFPT (magenta).

**Movie S2.** Contact inhibition of locomotion and interpenetration of WT cells **(A)** or α-catenin KO cells **(B), related to figure 1I**, expressing LifeAct-GFP (green) or LifeAct-TagRFPT (magenta).

**Movie S3.** Parallel actin flow at side-side junctions between leader cells expressing LifeAct-GFP (green) or LifeAct-TagRFPT (magenta) **(A), related to figure 2A**, and the effect on junctional E-cadherin (green) **(B), related to figure 2B**.

**Movie S4, related to Figure 2C**. Free edge of leader cells expressing LifeAct-TagRFPT (blue) and E-cadherin-GFPTG (green). Surface E-cadherin is labelled with surface-delivered E-cadherin mAbs (magenta). Yellow lines: E-cadherin-containing intracellular vesicles; red lines indicate: E-cadherin surface clusters.

**Movie S5, related to Figure 3F**. contact formation of WT cells expressing LifeAct-iRFP670 (blue), and α-catenin KO cells expressing LifeAct-TagRFPT (magenta) and E-cadherin-GFP^TG^ (green). Red arrow heads: anterograde movement of E-cadherin clusters.

**Movie S6, related to Figure 6C**.ss Lamellipodia dynamics and actin flows during contact formation of cells expressing LifeAct-GFP (green) or LifeAct-TagRFPT (magenta), and different α-catenin mutants.

**Movie S7, related to Figure 6F**. Contact inhibition of locomotion and interpenetration of cells expressing LifeAct-GFP (green) or LifeAct-TagRFPT (magenta), and different α-catenin mutants.

