## Supplemental material for "An E-cadherin-actin clutch translates the mechanical force of cortical flow for cell-cell contact to inhibit epithelial cell locomotion"

**This file includes:**

Computational Supplement
Tables S1 and S2
Figures S1 to S6

**Other Supplementary Materials for this manuscript include the following:**

Movies S1 to S7

### COMPUTATIONAL SUPPLEMENT

#### Introduction

To model the dynamics of formation of cadherin-mediated cell-cell adhesions, we adapted a previously described molecular clutch model [1], originally designed to simulate not cell-cell but cell-extracellular matrix adhesions.

In this adapted version, the model considers two cells that are interconnected via cadherin *trans*-interaction, which are in turn linked to the actin cytoskeleton. These cells will try to move away from one another, thereby loading a certain amount of force in the links that connect them.

#### Elements and dynamics of the system

The system consists of two cells (hereafter denoted as Right Cell (RC) and Left Cell (LC)) interconnected through a set of links, or "clutches". The movement of each cell will be represented as a filament of actin (F-actin), and filaments in both cells move in opposite directions to each other (antiparallel flows). These actin filaments will be bound to a series of adaptor proteins ($\alpha$-catenin and $\beta$-catenin) that, in turn, will bind to E-cadherin. Each complex (clutch) between the different proteins is a Cadherin-Catenin Complex (CCC). Cadherin trans-interaction enables the binding of each CCC to an equivalent CCC in the neighboring cell. Once CCCs bind to each other and to actin filaments in each cell, they resist antiparallel actin flows (Fig S6A).

The actin filament is pulled by $n_m$ myosin motors, each of which can exert a force $f_m$. When not submitted to any force, the filaments will move away from one another with a certain unloaded velocity $v_u$ and $-v_u$ for RC and LC respectively. Making use of experimental data, we consider that CCCs on both cells are precoupled to each other. Thus, the only binding/unbinding event that is explicitly considered by the model is the bond between each CCC and actin.

$k_{on}$ is the binding rate, that is the product of a true binding rate $k_{on_t}$times the density of CCC complexes available $d_{ccc}$ (that is directly related to the cadherin density, since we consider the whole complex to be precoupled): $k_{on} = k_{on_t} \cdot d_{CCC}$ .

Once CCCs are bound to actin on both cells, the clutch is considered to be engaged, and transmits force as a spring with spring constant $k_c$. CCCs are also allowed to unbind according to an unbinding rate $k_{off}$ This unbinding rate is assumed to depend on force as a two-state catch bond, as previously measured in $\alpha$-catenin-F-actin bonds [2]. These two states refer to a low affinity and a high affinity state, which depend on a conformational change of $\alpha$1-helix H1 (unfolding) that increases the affinity of $\alpha$-catenin for actin and increases the strength of the bond. As in any catch bond (or more precisely, catch-slip bond) increasing forces on CCCs first stabilize bonds up to a threshold force, and then weaken them when force rises above this threshold.

Apart from the engaging – disengaging feature of the links, we also model the process of adhesion growth. To this end, we assume that, as previously described, the unfolding of the $\alpha$1-helix H1 of the actin binding domain (ABD) of α-catenin (driven by force, see below) not only induces the transition between the weak and strong bound state (increasing the lifetime of the bond), but also promotes actin bundling (via dimerization) which further contributes to the growth of the AJ's [3]. Thus, we model helix unfolding through an unfolding rate $k_{unf}$. Once unfolded, we consider the helix bundling actin immediately. This actin bundling is assumed to increase the number of available links (clutches), which is modelled by increasing $d_{ccc}$ by an amount $d_{add}$. Of note, however, the model is not restricted to a bundling-mediated mechanism but is consistent with any mechanism by which ABD unfolding increases cadherin recruitment. If a clutch disengages from actin before this bundling can occur, $d_{ccc}$ decreases instead by an amount $d_{off}$, accounting for the fact that the complex is removed from the site where adhesion is happening and is no longer available (Fig S6B).

To summarize, $\alpha$-catenin is the protein that regulates the clutch behavior (and thereby force transmission), engaging with a certain on-rate depending on its binding affinity, and disengaging as a catch bond as a function of force. Force also induces a conformational change in its ABD increasing its effective on-rate via ABD dimerization and therefore actin bundling [4]. As a whole, $\alpha$-catenin acts as the mechanosensitive component on each of the protein chains that connect the moving actin filaments.

Time evolution of the system

Initially, we consider that both CCCs on each cell are interconnected via cadherin trans-interaction. At each timestep, the model evaluates the time required for all the different potential events to occur. This is done stochastically according to their corresponding rates, as described previously [5] (Fig S6C).

At each time step $t_s$:

- The $n_m$ myosin motors on RC and LC pull their actin filament (each one capable of exerting a maximum force $|f_m|$ ) in opposite directions at a velocity that depends on the total force transmitted $F(t)$:

$$v_{R,L}\ (t) = \pm v_u \left(1 - \frac{F(t)}{n_m \cdot f_m}\right)$$

  At the beginning of the simulation $F(0) = 0$, and the velocity coincides with the unloaded velocity $v_u$ for both sides. The complete stalling of the flow would happen when the force loaded on the system equals $F_{stall} = n_m \cdot f_m$.

  In each of the links, the connected clutches (CCC) will move away from one another, their starting point being $x_0 = 0$ in case they were disengaged. The one in the right will move following $x_R^{t+1} = x_R^t + v_R \cdot t_s > 0$ due to the right filament's velocity $v_R > 0$, and the one in the left $x_L^{t+1} = x_L^t + v_L \cdot t_s < 0$ due to the left filament's velocity $v_L < 0$[3]. The filaments' movement elongates each of the connected (engaged) springs a total quantity $\Delta x_c = x_R + v_R \cdot t_s - x_L - v_L \cdot t_s$.
- The elongation induces a total force $F = \sum_{i=1}^{n_e} k_c \Delta x_c$, where $n_e$ corresponds to the number of links simultaneously engaged. Note that the larger the load force is, the lower actin filament velocity will be.
- For each unbound CCC clutch, $\alpha$-catenin is allowed to bind to F-actin according to $k_{on}$. This binding rate establishes the connection between the CCC and the myosin-actin cytoskeleton. If one of the links becomes bound in both sides, that spring will become engaged.
- For the CCC clutches that are engaged, force exerted by the pulling of myosin motors from both sides can also unbind them with a $k_{off}$ rate that depends on force. If a certain clutch unbinds successfully, its position will be reset to $x_0 = 0$. Even if the

second clutch of the link is still bound, since there is now no real elongation of the spring its position will also be set to $x_0 = 0$. This rate is also different for clutches with folded or unfolded alpha 1 helices, as described below.

- Engaged links are also allowed to unfold the alpha 1 helix, leading to actin bundling. If this has occurred when the link unbinds, $d_{ccc}$ is increased by $d_{add}$. Otherwise, dcc is decreased by $d_{off}$.

At each time step the times corresponding to all possible events (binding, unfolding, unbinding) for each clutch are calculated. Then, only the events occurring before the duration of the time step $t_s$ are executed. As a control, we also carried out simulations using a Gillespie algorithm, in which only the event with the shortest time is executed, leading to time steps with a variable duration. The same trends were obtained after minor adjustments of parameters.

The system will eventually arrive to the stationary state that depends on the CCC density accumulated during the interval of time specified. Two distinct situations arise:

- CCC clusters are large enough to exert a force that enables both cortical flows to stop (at least, slow down to orders of magnitude smaller than the initial value). Stable AJ appears.
- The number of simultaneously engaged springs is not enough in order to build a force that would make the flow stop. In this situation the actin filaments would move away from one another. Stable AJ cannot appear.

Discussion of model parameters and assumptions

As regards the dynamics of the actomyosin contraction, the force exerted per motor is of the order of magnitude of that reported earlier [6]. However, in reality, the velocity of the filament depends upon two factors: (1) actin polymerization and depolymerization at the leading and trail edges and (2) myosin contraction. Although only myosin contractility is explicitly considered here, we note that this parameter can be considered to include both contributions, which would lead to the same effects in the model. The F-actin velocity is also in accordance with experiments, with a value $\approx 1\ \mu$m/min. The value for the clutch spring constant $k_c$ corresponds to the stiffness of the $\alpha$-catenin-actin link, which we assume to be high so as to load force very quickly. It must be noted, however, that this is not strictly required, and could be avoided by introducing instead a higher number of added cadherin per dimerization event. Indeed, a lower $k_c$would imply that more CCC-CCC links need to be simultaneously engaged to completely stop the flow. We decided not to include a dimerization rate variable $k_{dim}$, since α-catenin ABD unfolding promoted actin bundling in most of the cases, and simulations with or without it gave matching results. Except from $d_{add}$ and $k_{on}$, all model parameters were kept constant. All parameters and the values used in simulations are shown in Table S1. Importantly, an increase in both $d_{add}$ and $k_{on}$ enabled us to emulate the behavior of the $\alpha$ -catenin A+ mutant. We increased $k_{on}$ because of the exposed actin binding residues that increased the affinity of $\alpha$-catenin for actin, and $d_{add}$ was increased in comparison to its initial value as well, to account for a higher probability of dimerization. Together, tuning these two terms was sufficient to reproduce the results obtained in Fig 5I. $d_{off}$ was not modified, since the term corresponds to the CCC density that is lost on the cell surface due to rearward flow.

Finally, we considered the weakest link in the whole protein chain to be the bond between $\alpha$-catenin and actin. This assumption was in accordance with experiments in the fact that introducing a higher affinity $\alpha$-catenin mutant induced a higher surface cadherin flow on the free surfaces. However, we note that the bond lifetimes of some of the cadherin trans interactions are lower than the ones corresponding to our assumed weakest bond [7]. These bonds could be included in the model as another event leading to clutch disengagement. However, this would merely increase overall unbinding rates, something that could be compensated by tuning other model parameters (such as cadherin densities). Further, this would not predict any differences between the different conditions analyzed here. Thus, for the sake of simplicity, this was not done.

Two state catch bond model

We modeled the $\alpha$-catenin-F-actin interaction as a two state catch bond as described previously [2]. This model considers three states, unbound (state 0), a low affinity bound state (state 1), and a high affinity bound state (state 2). Thus, the dissociation rate from state 1 $k_{10}$ is much higher than the dissociation from state 2 $k_{20}$. Further, the transition rate $k_{12}$ (from state 1 to state 2) increases with force (slip pathway), whereas $k_{21}$ decreases with force (catch pathway). In the model presented here, this transition rate is due to the unfolding of the H1 in $\alpha$-catenin. Thus, state 0 is unbound, state 1 is bound-folded, and state 2 is bound-unfolded. Each of the expressions for the transition rates follow the Bell equation, which reads as follows:

$$k_{ij} = k_{ij}^{0} \cdot \exp\left(\mathrm{Fx_{ij}}/\mathrm{k_B T}\right)$$

Where $k_{ij}$ is the rate constant, $F$ the force exerted, $k_B$ the Boltzmann constant and T the absolute temperature. $k_{10}$ and $k_{20}$ correspond to unbinding rates for the folded and unfolded states, respectively. $k_{12}$ and $k_{21}$ correspond to the unfolding and refolding rates for the $\alpha_1$ helix, respectively.

The values used for the different $k_{ij}$, as well as its corresponding $k_{ij}^{0}$ in the formula, were taken from the aforementioned [2]. According to this, and thanks to the fact that $k_{12} \gg k_{21}$ we expected a great increase in the number of unfolded $\alpha$-catenin in the chains in comparison to the ones that remained folded (or went back to that state), thus further promoting cadherin accumulation at the cell junction.

| Parameter | Symbol | Values (MC) | Values (Gillespie) | Adjusted/Measured |
|---|---|---|---|---|
| Number of myosin motors | $n_m$ | 600 | 600 | Adjusted |
| Force exerted per motor | $f_m$ | 2 $pN$ | 2 $pN$ | Measured *(34)* |
| Unloaded velocity | $v_u$ | 15 $nm/s$ | 15 $nm/s$ | Measured |
| Number of clutches (one side) | $n_c$ | 600 | 600 | Adjusted |
| Cadherin density (CCC density) | $d_{cad} = d_{CCC}$ | 2000 $/\mu m^2$ | 2000 $/\mu m^2$ | Adjusted |
| True on-rate | $k_{on}$ | $1.3 \times 10^{-4}$ $\mu m^2/s$ (WT)<br>$2.8 \times 10^{-4}$ $\mu m^2/s$ (A+) | $3.2 \times 10^{-4}$ $\mu m^2/s$ (WT)<br>$3.3 \times 10^{-4}$ $\mu m^2/s$ (A+) | Adjusted |
| Spring constant | $k_c$ | $1.5 \times 10^{-2}$ $N/m$ | $1.5 \times 10^{-2}$ $N/m$ | Adjusted |
| Added cadherin density | $d_{add}$ | 6.4 $/(s \cdot \mu m^2)$ (WT)<br>13.2 $/(s \cdot \mu m^2)$ (A+) | 17.2 $/(s \cdot \mu m^2)$ (WT)<br>18.4 $/(s \cdot \mu m^2)$ (A+) | Adjusted |
| Substracted cadherin density | $d_{off}$ | 0.25 $/(s \cdot \mu m^2)$ | 0.32 $/(s \cdot \mu m^2)$ | Adjusted |

**Table S1. Model parameters.** For $\mathrm{d_{add}}$ and $\mathrm{k_{on}}$, both values for WT and A+ mutant are shown.

| Transition | Symbol | Value |
|---|---|---|
| 1→0 | $k_{10}^{0}$ | $11/s$ |
| | $x_{10}$ | 0.0 $m$ |
| 1→2 | $k_{12}^{0}$ | $3/s$ |
| | $x_{12}$ | 0.2x10$^{-9}$ $m$ |
| 2→1 | $k_{21}^{0}$ | $20/s$ |
| | $x_{21}$ | -4x10$^{-9}$ $m$ |
| 2→0 | $k_{20}^{0}$ | $0.14/s$ |
| | $x_{20}$ | 0.4x10$^{-9}$ $m$ |

**Table S2. Parameters values in the Bell equation for the different transition rates in the two state catch bond model.** Model parameters extracted from [2].

### SUPPLEMENTAL FIGURES

Figure S1 Noordstra *et al.*

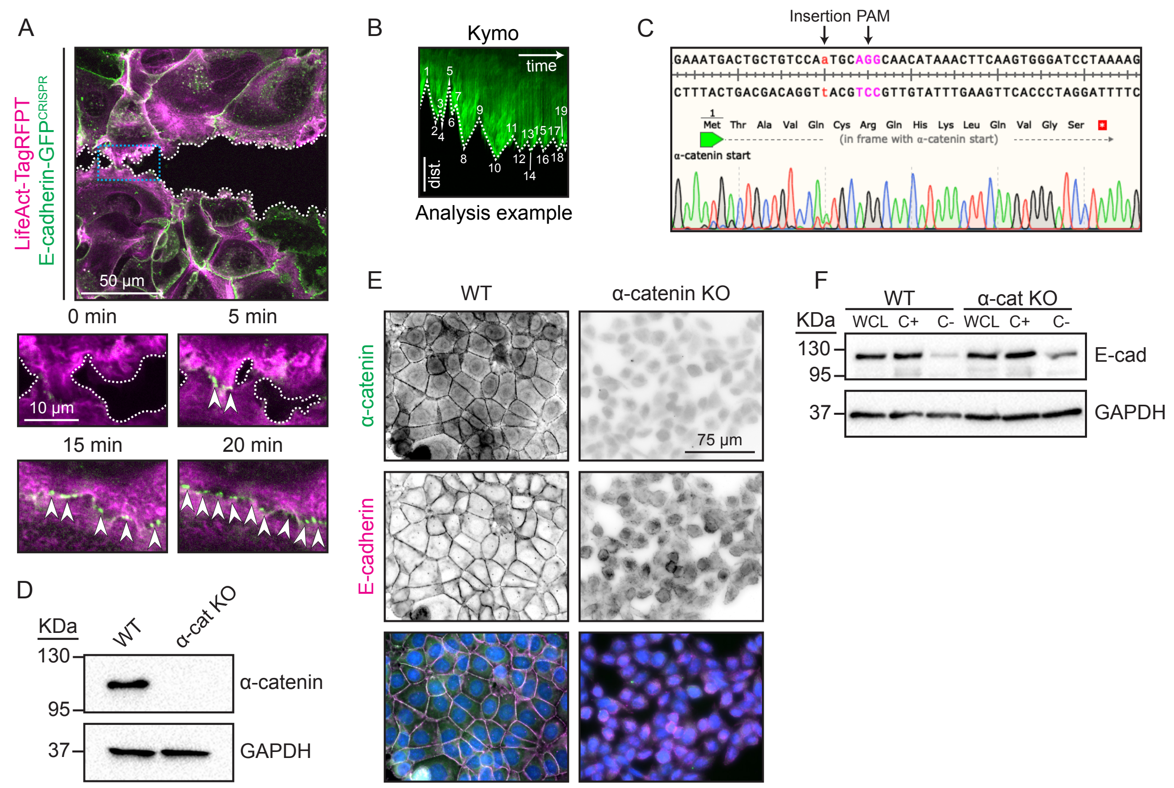


**Figure S1.**

**(A)** Junction formation between MCF7 cells expressing LifeAct-TagRFPT (magenta) and E-cadherin-GFP$^{CRISPR}$ (green). White dotted line: free cell edge; arrow heads: Adherens Junctions (AJ).

**(B)** Analysis example lamellipodia dynamics. Every directional change in the kymograph (switch from protrusive to retractive motion or vice versa) is counted as 1.

**(C)** Genomic sequencing results of CTNNA1, exon 1, in $\alpha$-catenin KO cells.

**(D)** Western blot analysis of MCF7 WT and $\alpha$-catenin KO cells.

**(E)** Confluent layer of WT and $\alpha$-catenin KO cells stained for $\alpha$-catenin (green) and E-cadherin (magenta).

**(F)** Trypsin protection assay and western blot analysis of WT and $\alpha$-catenin KO cells. Surface E-cadherin is protected against trypsin cleavage by the addition of $Cacl_2$. WCL = Whole Cell Lysate, C+ = with $Cacl_2$ (Total E-cadherin levels), C- = without $Cacl_2$ (Intracellular E-cadherin only).

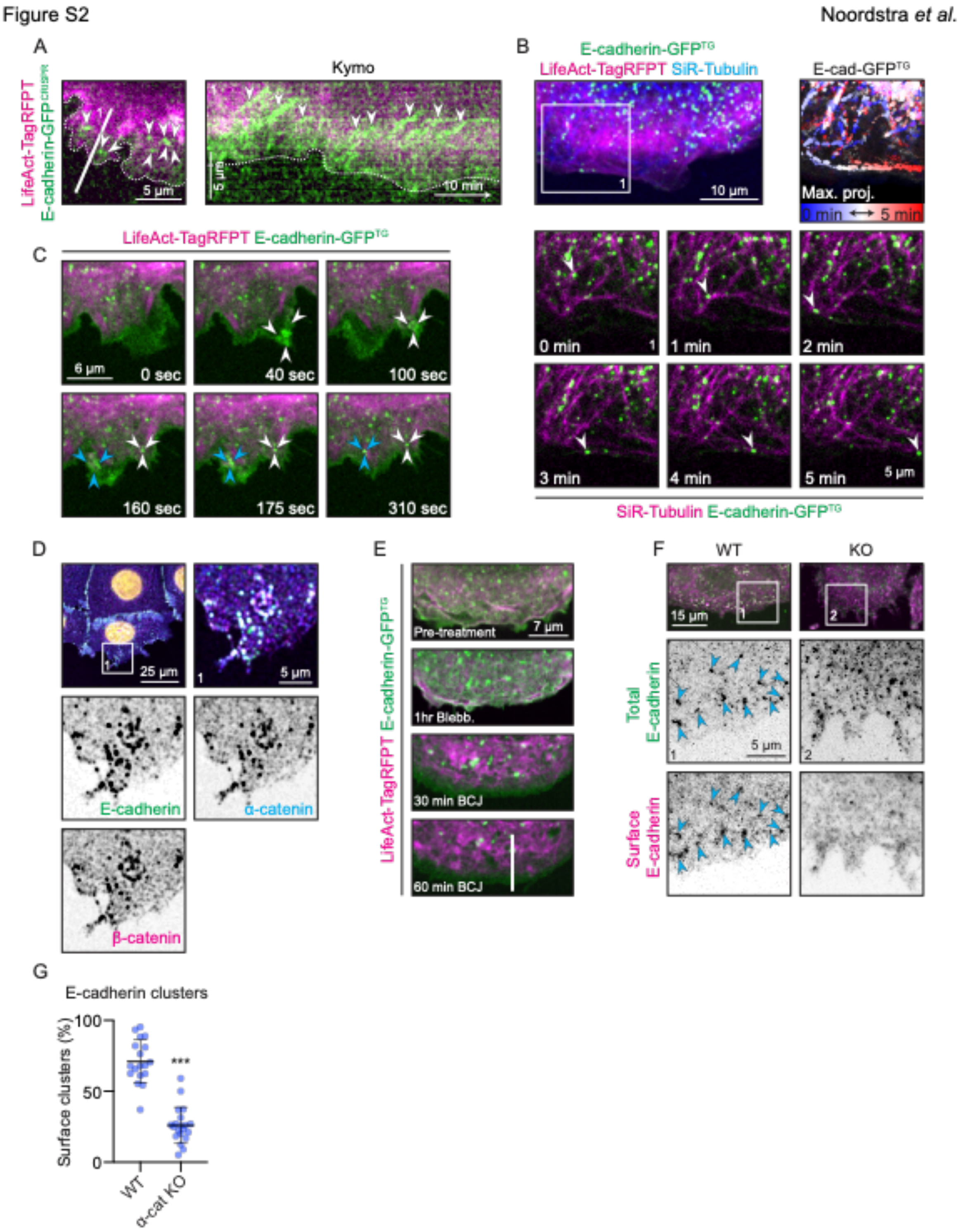


**Figure S2.**

**(A)** MCF7 leader cells expressing LifeAct-TagRFPT (magenta) and E-cadherin-GFPCRISPR (green). Wavy white dotted line: free cell edge; straight white dotted line: site of kymograph; white arrow heads: retrograde movement (Kymo) of E-cadherin clusters.

**(B)** Leader cells expressing E-cadherin-GFP$^{TG}$ (green), LifeAct-TagRFPT (magenta), stained for microtubules using SiR-Tubulin (blue in overview, magenta in zooms). White arrow heads: E-cadherin vesicle moving over a microtubule.

**(C)** Leader cells expressing LifeAct-TagRFPT (magenta) and E-cadherin-GFP$^{TG}$ (green). White and blue arrow heads: E-cadherin cluster formation in the lamellipodia.

**(D)** Leader cells stained for E-cadherin (green), β-catenin (magenta) and α-catenin (blue).

**(E)** Leader cells expressing LifeAct-TagRFPT (magenta) and E-cadherin-GFP$^{TG}$ (green). Images show effect of BCJ treatment on E-cadherin and actin. White dotted line: site of kymograph (2E).

**(F, G)** WT and α-catenin KO leader cells stained for total E-cadherin (green) and surface E-cadherin (magenta). (F) Representative images. Blue arrow heads: E-cadherin surface clusters. (G) Quantification of E-cadherin surface clusters. WT cells (n=18), α-catenin KO cells (n=20) from 2 independent experiments.

***P<0,001; Mann-Whitney U test. Data are means ±SD with individual datapoints indicated.

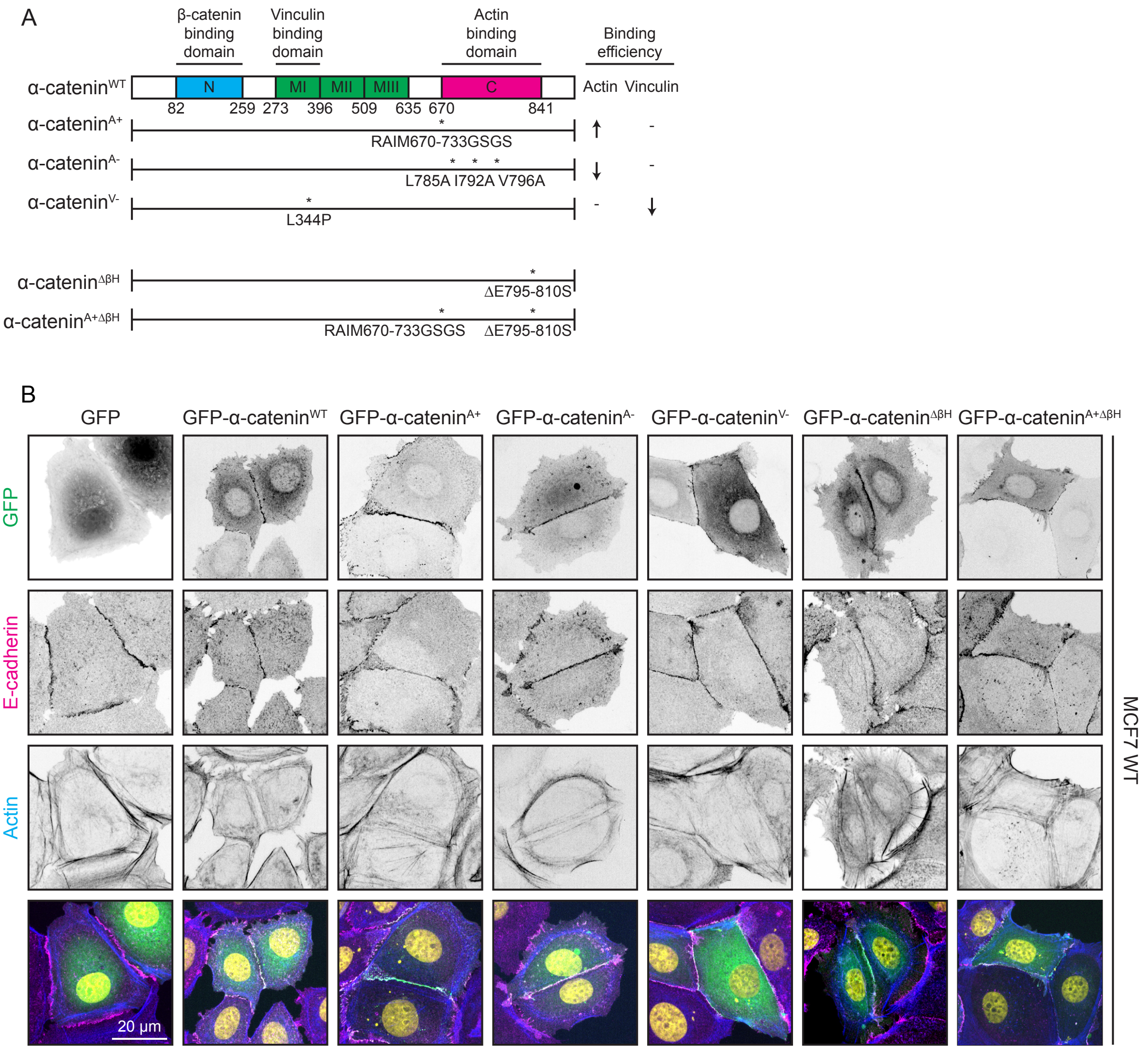


**Figure S3.**

**(A)** Domain organization, mutation indications and binding efficiencies of α-catenin$^{WT}$ and all α-catenin mutants used in this study.

**(B)** Localization of GFP-α-catenin transgenes (green) in WT cells co-stained for E-cadherin (magenta) and actin (blue). All α-catenin mutants are properly incorporated into adherens junctions, indicating β-catenin binding is not affected.

A

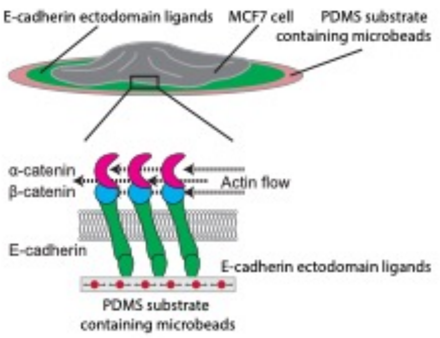


**Figure S4.**

**(A)** Schematic representation of a cell spread on an E-cadherin ectodomain-coated PDMS substrate and the effect of actin flow on the cadherin-catenin complexes.

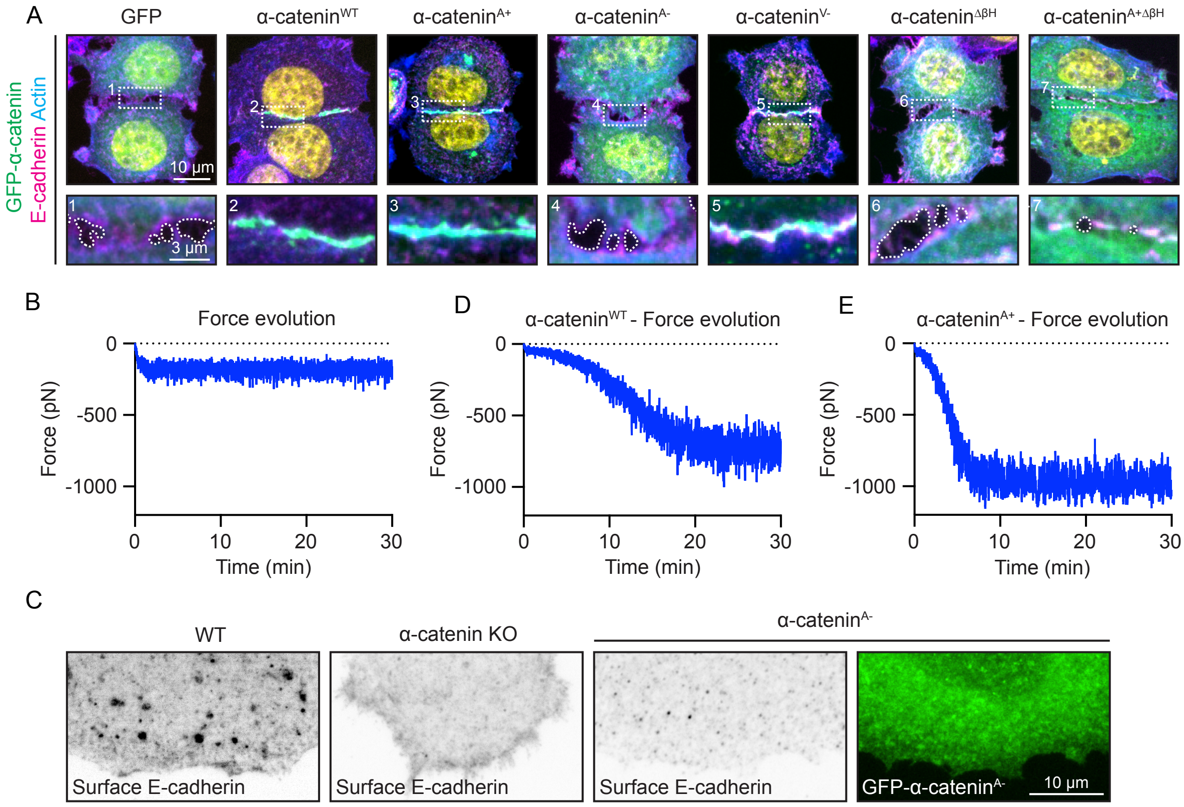


**Figure S5.**

**(A)** MCF7 α-catenin KO cells expressing GFP, GFP-α-catenin$^{WT}$, GFP-α-catenin$^{A+}$, GFP-α-catenin$^{A-}$, GFP-α-catenin$^{V-}$, GFP-α-catenin$^{\Delta\beta H}$ or GFP-α-catenin$^{A+\Delta\beta H}$ (green), stained for E-cadherin (magenta) and actin (blue). White dotted line: free cell edge.

**(B)** Force evolution corresponding to (5C)

**(C)** Surface E-cadherin clusters in leader WT cell, α-catenin KO cell and α-catenin KO cell reconstituted with GFP-α-catenin$^{A-}$ (green).

**(D)** Force evolution corresponding to (6A)

**(E)** Force evolution corresponding to (6B)

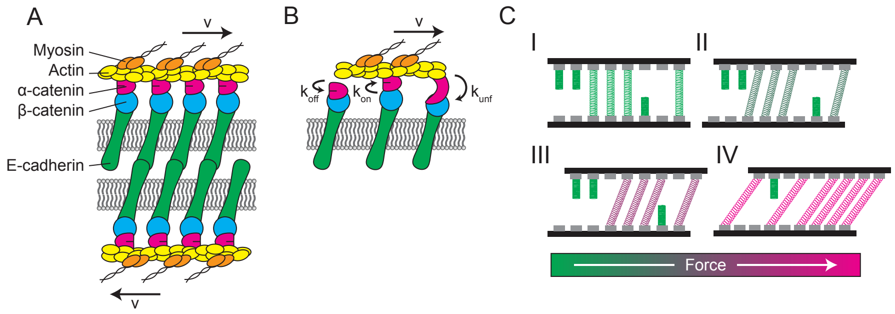


**Figure S6.**

**(A)** Scheme of the molecular ensemble. Actin filaments of neighboring cells are connected via a set of CCC-CCC links. CCC from one cell bind through cadherin trans interaction to the equivalent CCC from the other cell. α-catenin-F-actin bond regulates force transmission between actin filaments throughout the whole link.

**(B)** Dynamics of the clutch. As regards F-actin – $\alpha$-catenin bond, it can either unbind with a $k_{off}(F)$, bind according to $k_{on}(d_{CCC})$ or unfold following $k_{unf}(F)$.

**(C)** Time evolution of the system. I Initial random distribution of clutches, some engaged and others not. II As time passes, actin filaments begin to move in opposite directions, thereby loading some force on the system due to the tension induced on the springs. III As the actin filaments continue to pull, deformation on the spring becomes greater, and so it does the total loaded force, decreasing the actin filament movement. IV High tension on the links induces an increase on the number of clutches in the system, thereby generating a loaded force able to completely stop the flow.
