## Supplementary material for "An E-cadherin-actin clutch translates the mechanical force of cortical flow for cell-cell contact to inhibit epithelial cell locomotion": Key recources table

| REAGENT or RESOURCE | SOURCE | IDENTIFIER |
|---|---|---|
| **Antibodies** | | |
| Mouse monoclonal anti E-cadherin ectodomain | BD Biosciences | #563571 |
| Mouse monoclonal anti β-catenin | BD Biosciences | #610154 |
| Mouse monoclonal anti E-cadherin | Gift from Dr. M. Takeichi | N/A |
| Rabbit polyclonal anti GAPDH | R&D Systems | #2275-PC-100 |
| Rabbit polyclonal anti α-catenin | Thermo Fisher | #71-1200 |
| Rabbit polyclonal anti α-catenin | Gift from Dr. B. M. Gumbiner | N/A |
| Rabbit polyclonal anti α-catenin (VD7) | Vestweber lab | N/A |
| Rat monoclonal anti E-cadherin | Abcam | #ab11512 |
| Rat monoclonal anti E-cadherin | Thermo Fisher | #13-1900 |
| Rat monoclonal anti α-catenin (α18) | Gift from Dr. A. Nagafuchi | N/A |
| Goat anti mouse, rabbit, rat Alexa-Fluor-405 | Thermo Fisher | #A-31553<br>#A-48254<br>#A-48261 |
| Goat anti mouse, rabbit, rat Alexa-Fluor-488 | Thermo Fisher | #A-11001<br>#A-11008<br>#A-11006 |
| Goat anti mouse, rabbit, rat Alexa-Fluor-594 | Thermo Fisher | #A-11032<br>#A-11037<br>#A-48264 |
| Goat anti mouse, rabbit, rat Alexa-Fluor-647 | Thermo Fisher | #A-21236<br>#A-21245<br>#A-48265 |
| Goat anti mouse HRP | Bio-Rad | #1706516 |
| Goat anti rabbit HRP | Bio-Rad | #1706515 |
| **Bacterial and virus strains** | | |
| NEB 10-beta Competent *E.coli* | New England Biolabs | #C3019H |
| Stellar Competent *E.coli* | Clontech | #636766 |
| **Chemicals, peptides, and recombinant proteins** | | |
| Lipofectamin 3000 | Thermo Fisher | #L3000015 |
| Puromycin | Sigma-Aldrich | #P8833 |
| G418 | Santa Cruz | #108321-42-2 |
| Lenti-X concentrator | Clontech | #631232 |
| Alexa Fluor® 488, 594, 647 Phalloidin | Thermo Fisher | #A12379<br>#A12381<br>#A22287 |
| SiR-Tubulin | Cytoskeleton | #CY-SC002 |
| Para-nitroblebbistatin | Optopharma | #DR-N-111 |
| CK666 | Sigma-Aldrich | #SML0006 |
| Jasplakinolide | Merck | #420107 |

| | | |
|---|---|---|
| Protein A | Thermo Fisher | #21181 |
| hE-cadherin/Fc | SinoBiological | #10204 |
| CY-52-276 Part A | Dow Corning Toray | #CY 52-276 A |
| CY-52-276 Part B | Dow Corning Toray | #CY 52-276 A |
| 3-aminopropyl trimethoxysilane | Sigma Aldrich | #281778 |
| Crystalline Trypsin | Sigma Aldrich | #T-0303 |
| Chemiluminescent Substrate | Thermo Fisher | #34579 |
| ProLong Gold with DAPI | Cell Signaling | #8961 |
| ProLong Gold without DAPI | Cell Signaling | #9071 |
| **Experimental models: Cell lines** | | |
| MCF7 | ATCC | HTB-22 |
| HEK293T | ATCC | CRL-3216 |
| MCF7 LifeAct-TagRFPT | This study | N/A |
| MCF7 LifeAct-TagRFPT, E-cadherin-GFP$^{CRISPR}$ | This study | N/A |
| MCF7 LifeAct-TagRFPT, E-cadherin-GFP$^{TG}$ | This study | N/A |
| MCF7 LifeAct-GFP | This study | N/A |
| MCF7 $\alpha$-catenin KO, LifeAct-TagRFPT | This study | N/A |
| MCF7 $\alpha$-catenin KO, LifeAct-GFP | This study | N/A |
| MCF7 $\alpha$-catenin KO, LifeAct-TagRFPT, E-cadherin-GFP$^{TG}$ | This study | N/A |
| MCF7 LifeAct-iRFP670 | This study | N/A |
| MCF7 LifeAct-TagRFPT, iRFP670-$\alpha$-catenin$^{WT}$ | This study | N/A |
| MCF7 LifeAct-GFP, iRFP670-$\alpha$-catenin$^{WT}$ | This study | N/A |
| MCF7 LifeAct-TagRFPT, iRFP670-$\alpha$-catenin$^{A+}$ | This study | N/A |
| MCF7 LifeAct-GFP, iRFP670-$\alpha$-catenin$^{A+}$ | This study | N/A |
| MCF7 LifeAct-TagRFPT, iRFP670-$\alpha$-catenin$^{\Delta\beta H}$ | This study | N/A |
| MCF7 LifeAct-GFP, iRFP670-$\alpha$-catenin$^{\Delta\beta H}$ | This study | N/A |
| MCF7 LifeAct-TagRFPT, iRFP670-$\alpha$-catenin$^{A+\Delta\beta H}$ | This study | N/A |
| MCF7 LifeAct-GFP, iRFP670-$\alpha$-catenin$^{A+\Delta\beta H}$ | This study | N/A |
| MCF7 LifeAct-TagRFPT, E-cadherin-GFP$^{TG}$, iRFP670-$\alpha$-catenin$^{WT}$ | This study | N/A |
| MCF7 LifeAct-TagRFPT, E-cadherin-GFP$^{TG}$, iRFP670-$\alpha$-catenin$^{A+}$ | This study | N/A |
| MCF7 LifeAct-TagRFPT, E-cadherin-GFP$^{TG}$, iRFP670-$\alpha$-catenin$^{\Delta\beta H}$ | This study | N/A |
| MCF7 LifeAct-TagRFPT, E-cadherin-GFP$^{TG}$, iRFP670-$\alpha$-catenin$^{A+\Delta\beta H}$ | This study | N/A |
| **Oligonucleotides** | | |
| *CTNNA1* targeting sequence 5’GAAATGACTGCTGTCCATGC‘3 | This study | N/A |
| **Recombinant DNA** | | |
| PX459 CTNNA1 ($\alpha$-cateninWT CRISPR KO) | This study | N/A |
| PLViP-LifeAct-GFP | This study | N/A |
| PLViP-LifeAct-TagRFPT | This study | N/A |

| | | |
|---|---|---|
| PLViP-LifeAct-iRFP670 | Gift from Dr. J van Buul | N/A |
| pLViN-E-cadherin-GFP$^{TG}$ | This study | N/A |
| GFP-$\alpha$-catenin$^{WT}$ | This study | N/A |
| GFP-$\alpha$-catenin$^{A+}$ | This study | N/A |
| GFP-$\alpha$-catenin$^{A-}$ | This study | N/A |
| GFP-$\alpha$-catenin$^{V-}$ | This study | N/A |
| GFP-$\alpha$-catenin$^{\Delta\beta H}$ | This study | N/A |
| GFP-$\alpha$-catenin$^{A+\Delta\beta H}$ | This study | N/A |
| PLViN-iRFP670-$\alpha$-catenin$^{WT}$ | This study | N/A |
| PLViN-iRFP670-$\alpha$-catenin$^{A+}$ | This study | N/A |
| PLViN-iRFP670-$\alpha$-catenin$^{\Delta\beta H}$ | This study | N/A |
| PLViN-iRFP670-$\alpha$-catenin$^{A+\Delta\beta H}$ | This study | N/A |
| **Software and algorithms** | | |
| ImageJ 1.52 | | https://imagej.nih.gov/ij/download.html |
| Prism 9.0.0 | Graphpad | https://www.graphpad.com/scientific-software/prism/ |
| Matlab 9.9 | Mathworks | https://www.mathworks.com/products/matlab.html |
| LASX | Leica | https://www.leica-microsystems.com/products/microscope-software/p/leica-las-x-ls/ |
| Fusion | Andor | https://fusion.help.andor.com/display/fusionum/Home |
| Zen Blue 3 | Zeiss | https://www.zeiss.com/microscopy/int/products/microscope-software/zen.html |
| **Other** | | |
| 4-well silicone inserts | Ibidi | #F8807 |
| Carboxylated fluorescent beads | Thermo Fisher | #80469 |
